# A disrupted compartment boundary underlies abnormal cardiac patterning and congenital heart defects

**DOI:** 10.1101/2024.02.05.578995

**Authors:** Irfan S. Kathiriya, Martin H. Dominguez, Kavitha S. Rao, Jonathon M. Muncie-Vasic, W. Patrick Devine, Kevin M. Hu, Swetansu K. Hota, Bayardo I. Garay, Diego Quintero, Piyush Goyal, Megan N. Matthews, Reuben Thomas, Tatyana Sukonnik, Dario Miguel-Perez, Sarah Winchester, Emily F. Brower, André Forjaz, Pei-Hsun Wu, Denis Wirtz, Ashley L. Kiemen, Benoit G. Bruneau

## Abstract

Failure of septation of the interventricular septum (IVS) is the most common congenital heart defect (CHD), but mechanisms for patterning the IVS are largely unknown. Here, we show that a *Tbx5^+^/Mef2cAHF*^+^ progenitor lineage forms a compartment boundary bisecting the IVS. This coordinated population originates at a first- and second heart field interface. Ablation of *Tbx5^+^/Mef2cAHF*^+^ progenitors cause IVS disorganization, right ventricular hypoplasia and mixing of IVS lineages. Reduced dosage of the CHD transcription factor TBX5 disrupts boundary position and integrity, resulting in ventricular septation defects (VSDs) and patterning defects, including misexpression of *Slit2* and *Ntn1*, which encode guidance cues. Reducing NTN1 dosage partly rescues cardiac defects in *Tbx5* mutant embryos. Loss of *Slit2* or *Ntn1* causes VSDs and perturbed septal lineage distributions. Thus, we identify *Tbx5* as a candidate selector gene, directing progenitors and regulating essential cues, to pattern a compartment boundary for proper cardiac septation, revealing mechanisms for cardiac birth defects.

## Main text

Organogenesis relies on fine tissue patterning, and disturbances cause organ malformations that result in birth defects. Examples of prevalent birth defects resulting from abnormal patterning include congenital limb abnormalities, neural tube defects or congenital heart defects (CHDs).

CHDs are the most common birth defects and are a leading cause of morbidity and mortality in childhood ^1^. CHDs are thought to result from alterations to the orchestrated patterning of heart development. Nearly half of all patients with CHDs have atrial septal defects (ASDs) or ventricular septal defects (VSDs), which is an abnormality of the formation of the interventricular septum (IVS) ^2,3^. Septal defects can occur either in isolation or combined with other anatomic defects, including abnormal chamber formation. For example, atrioventricular (AV) canal defects include VSDs, ASDs and abnormal development of the AV valves, with severe cases leading to chamber hypoplasia and functional single ventricle physiology ^4^. There is a large gap in our understanding of how early developmental events pattern cardiac morphogenesis and anatomy. Addressing this understudied need can inform diagnostic prognosis, family planning and therapeutic approaches for CHDs.

Complete atrioventricular septation into four chambers allows for separation of systemic and pulmonary circulations, and enables higher levels of oxygen for transport in arterial blood. Although formation of the IVS, which separates the left ventricle (LV) and right ventricle (RV), is an intricate and poorly understood process, some progress has been made to determine the embryonic origins of the IVS. Dye-labeling studies in chick have shown that the IVS is derived from the bulbo-ventricular region between the future RV and LV ^5^, and both the RV and LV supply compact layer cells and some lateral trabecular cells to form the core of the IVS ^6–8^. Likewise, discovery of the *Ganglion Nodosum* epitope from chick, which demarcates the primary interventricular foramen and subsequently the IVS, AV junction and ventricular conduction system in mouse, chick and humans, has led to the description of a “primary ring” for the ventricular septum ^9–11^. Consistent with the “primary ring”, cell behaviour of IVS primordia displays circumferential oriented cell growth from the interventricular groove at the outer curvature to the inner curvature at E10.5 ^12^. In addition, some gene expression domains are enriched in the IVS. For example, *Tbx5* is expressed in the LV and IVS, and proper TBX5 dosage is essential for IVS formation ^13^. *Irx2* is expressed in the IVS and adjacent ventricular tissue ^14^, and *Lyz2* is expressed at the bulbo-ventricular groove at E9.5 at the site of the future IVS ^15^. Further, the *Lyz2^+^* lineage contributes cells to the muscular portion throughout the developing IVS ^15^. This evidence delineates the embryonic origins of the IVS as a distinct region of the developing heart, although its earliest progenitors and regulation are unclear.

Studies have suggested that the linear heart tube is organized into specific segments ^16,17^, including the IVS ^5,18^. It is increasingly apparent in the developing mouse heart that discrete early cardiac progenitors in mesoderm are already fated to contribute to regionalization of specific cardiac anatomy ^19–26^. The first heart field (FHF) gives rise primarily to the left ventricle (LV) and parts of the atria, while the second heart field (SHF) contributes predominantly to the right ventricle (RV), outflow tract (OFT) and portions of the atria ^21^. The heart fields can be largely recapitulated by marking lineage-labelled progenitors at gastrulation ^24,25^. Specifically, a cell lineage labelled by the CHD-linked transcription factor *Tbx5* (*Tbx5*^+^ lineage) contributes to the LV, IVS and atria, while the anterior heart field enhancer of *Mef2c* (*Mef2cAHF*^+^) lineage largely contributes to the RV, IVS and OFT ^25^.

An aspect of proper tissue patterning entails the establishment of compartment boundaries to separate neighboring fields of cells that exhibit discrete functions ^27–29^. Few compartment boundaries have been identified in mammals, notably the dorso-ventral patterning of the limb ^30–32^ and the midbrain-hindbrain boundary ^33^. In the developing mouse heart, retrospective clonal cell analysis shows segregation of LV- and RV-lineages on either the left or right side of the IVS, establishing a presumptive mutual border between the two non-intermingling compartments ^12,25,34^, reminiscent of a compartment boundary^10–12^. Moreover, this evidence has led to a prediction of a compartment boundary at the IVS that acts as a barrier to cell mixing between the LV and RV ^12,25,34^. Using an intersectional-lineage labelling approach, a subset of cardiac progenitor cells labeled by both *Tbx5^+^/Mef2cAHF*^+^ were described with contributions to the left side of the IVS, delineating a cell lineage of an apparent compartment boundary ^25^. Understanding the roles of a *Tbx5^+^/Mef2cAHF*^+^ lineage-labeled compartment boundary at the IVS may reveal clues about IVS patterning with potential relevance for VSD etiologies. More broadly, how disturbances to compartment boundaries contribute to birth defects is largely unknown.

Here, we combined genetic lineage labeling with lightsheet microscopy to follow contributions of *Tbx5^+^/Mef2cAHF*^+^ precursors to a compartment boundary during cardiac morphogenesis. We ablated *Tbx5^+^/Mef2cAHF*^+^ precursors to determine the essential roles of the lineage-labeled compartment boundary for cardiac septation and chamber development. By leveraging a genetic lesion of the CHD gene *Tbx5,* our work uncovers the genetic regulation of the *Tbx5^+^/Mef2cAHF*^+^ lineage-labeled compartment boundary for proper IVS patterning. We deployed single cell RNA sequencing (scRNA-seq) to discover TBX5-sensitive cues that are essential for compartment boundary regulation and formation of the IVS during heart development.

### Early cardiac progenitors for the IVS, IAS and AV complex regions

We used conditional dual lineage labelling by *Tbx5^CreERT2/+^* or *Mef2cAHF-DreERT2* and marked cells during gastrulation at embryonic day (E) 6.5 by a single dose of tamoxifen-induced recombination. We followed separate reporter-labeled lineages by epifluorescence microscopy, histology or lightsheet microscopy (Fig. 1a-f). At E14.5, the *Tbx5*^+^ lineage contributed to the left side of the IVS, often ending at a presumptive line marked at the apex by the interventricular (IV) groove, consistent with previous reports suggesting a lineage boundary ^12,25,34^. *Tbx5*^+^ and *Mef2cAHF*^+^ lineages showed largely complementary patterns, with notable overlap of *Tbx5*^+^ and *Mef2cAHF*^+^ lineages at the IVS, AV complex region, and inter-atrial septum (IAS) (Fig. 1b, d, f).

**Figure 1.**
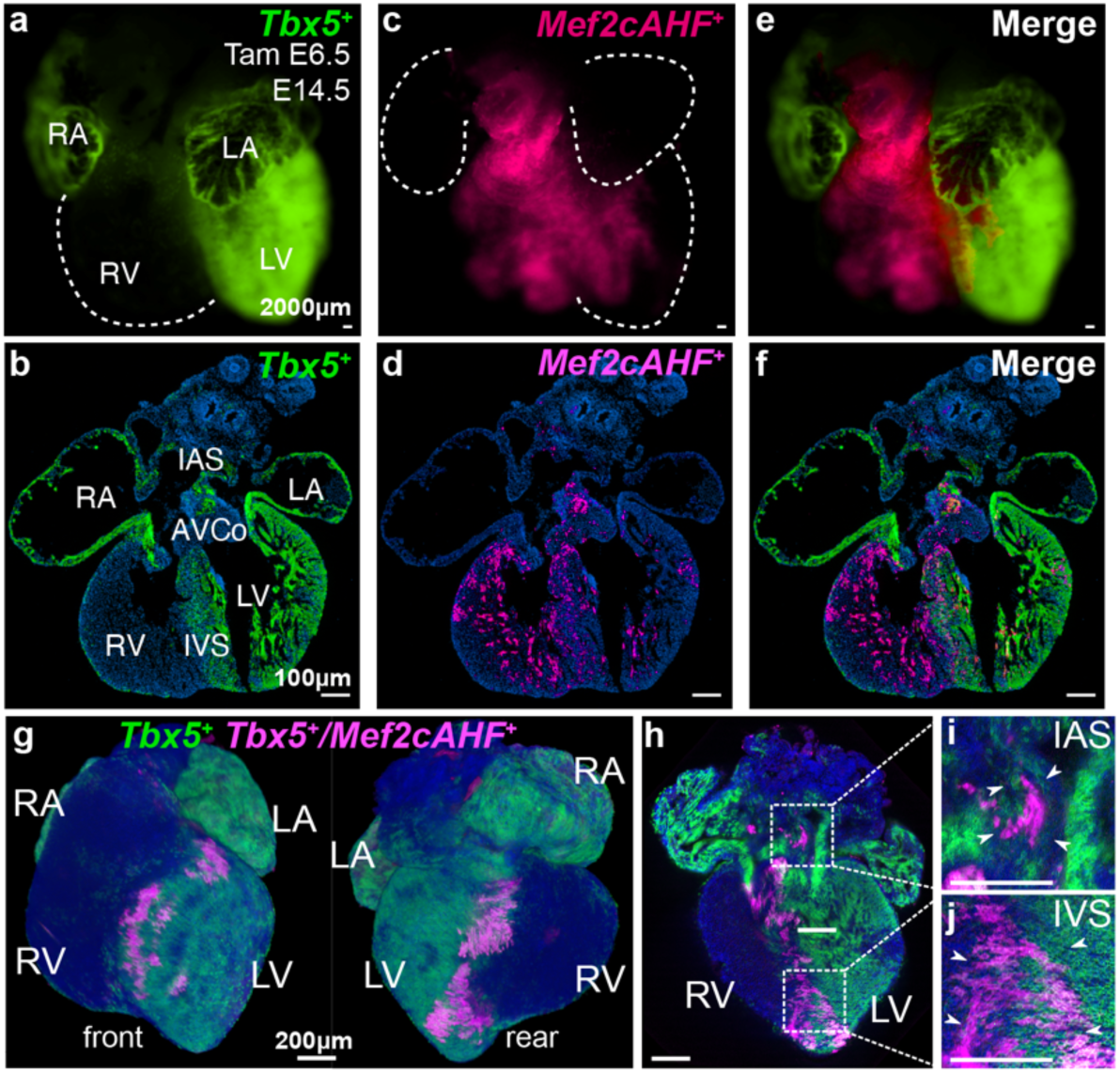
*Tbx5^+^/Mef2cAHF^+^* lineage marks a compartment boundary at the cardiac interventricular septum (IVS). **a**, **b**, At E14.5, clonal cell descendants of a *Tbx5^+^* lineage (ZsGreen) labeled at E6.5 contributes to the LV. **c**, **d**, a *Mef2cAHF^+^* lineage (tdTomato immunostaining) contributes to the RV, demonstrating largely complementary patterns. **e**, These lineages overlap (*Tbx5^+^/Mef2cAHF*^+^) at the IVS, as well as the atrioventricular complex (AVCo) region, and inter-atrial septum (IAS). *Tbx5^CreERT2^*^/+^;*Mef2cAHF-DreERT2*;*ROSA26^Ai6^*^/*Ai66b*^ hearts by epifluorescence microscopy (a, c, e; scale bars: 200 microns) and cryosections (b, d, f; scale bars: 100 microns). **g-j**, Maximal projection images by lightsheet microscopy of an intersectional reporter for the *Tbx5^+^/Mef2cAHF^+^* lineage (tdTomato immunostaining) in *Tbx5^CreERT2^*^/+^;*Mef2cAHF-DreERT2*;*ROSA26^Ai66^*^/+^ embryos at E14.5. Scale bars: 200 microns.

To better visualize the *Tbx5^+^/Mef2cAHF*^+^ lineage, we used a conditional intersectional-lineage labeling approach. Using a lineage reporter that is responsive to both CreERT2 and DreERT2 ^35^, we observed a spatial pattern of the *Tbx5^+^/Mef2cAHF*^+^ lineage in the IVS, AV complex region, and IAS (Fig. 1g). At E14.5, the septal lineage extended to the apex of the heart via the IVS, to the base of the heart at the AV complex region, as well as to the IAS superiorly^15–18^ (Fig. 1h-j). These results expanded upon previous findings in the developing IVS at E10.5 ^25^, by identifying contributions from *Tbx5^+^/Mef2cAHF*^+^ progenitors to additional anatomic sites during chamber formation and septation. Moreover, lineage contributions to sites of chamber septation were derived from *Tbx5^+^/Mef2cAHF*^+^ progenitors at E6.5 (Extended Data Fig. 1a-c), but not later at E8.5 or E10.5, implicating a narrow, early window for capturing *Tbx5^+^/Mef2cAHF*^+^ septal progenitors during cardiac mesoderm formation.

### *Tbx5^+^/Mef2cAHF*^+^ septal lineage is prefigured in early mouse heart development

To characterize the lineage derived from *Tbx5^+^/Mef2cAHF*^+^ septal progenitors during early heart development, we examined the localization of the lineage prior to heart tube formation at E8.0 (Extended Data Fig. 1d-f, Fig. 2, Video 1). By lightsheet imaging, the pattern of the *Tbx5^+^/Mef2cAHF*^+^ lineage was detected at the dorsal aspect of the *Tbx5*^+^ lineage, at an interface between *Tbx5*^+^ and *Tbx5*^-^ lineage compartments (Fig. 2a-e”). Cell segmentation at this early stage revealed that the *Tbx5^+^/Mef2cAHF*^+^ lineage resembled a nascent halo of cells that became readily apparent at later stages of heart development (Fig. 2f-i).

**Figure 2.**
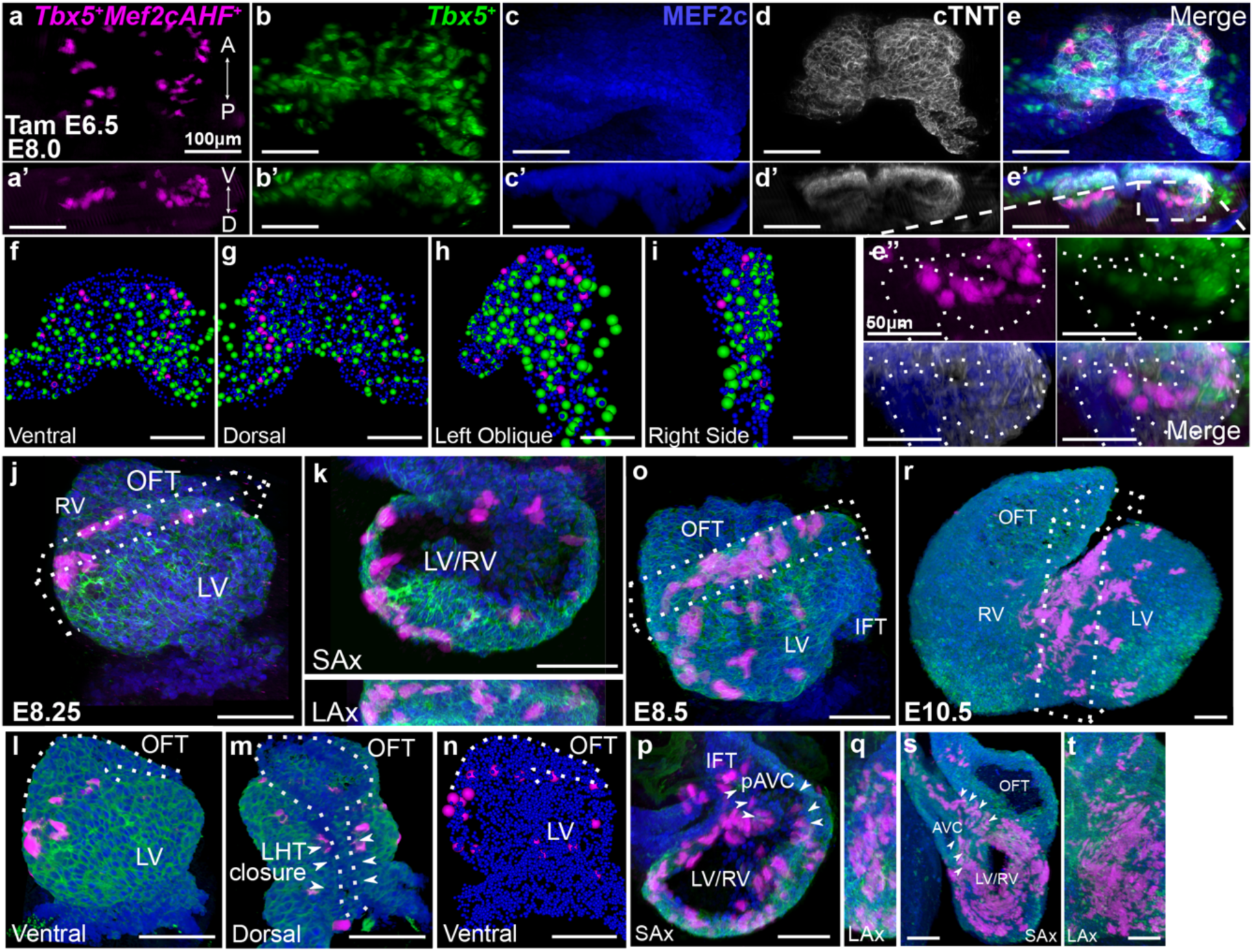
*Tbx5^+^/Mef2cAHF^+^* lineage is prefigured in early mouse heart development. **a-i**, Frontal (a-e), transverse (a’-e’), magnified transverse (e’’), and (f-i) cell segmentation views of maximum Z-projections of whole mount embryos by lightsheet imaging at E8.0 show the *Tbx5*^+^ lineage (ZsGreen), and immunostaining of tdTomato for the *Tbx5^+^/Mef2cAHF*^+^ lineage, MEF2c and cardiac troponin T (cTNT) in *Tbx5^CreERT2^*^/+^;*Mef2cAHF-DreERT2*;*ROSA26^Ai6^*^/*Ai66*^ embryos. A-P, anterior-posterior axis; V-D, ventral-dorsal axis. At E8.0 (approximately 3 somites), *Tbx5^+^* lineage cells are restricted to a ventral domain, whereas *Tbx5^+^/Mef2cAHF^+^* lineage cells originate along a boundary separating presumptive *Tbx5*^+^ and *Tbx5*^-^-lineage compartments. *Tbx5^+^/Mef2cAHF^+^* lineage cells lie in an apparent planar ringlet in the dorsal-ventral axis that expands as the heart tube grows. **j-n**, At E8.25 (approximately 6 somites), with dorsal closure of the linear heart tube (LHT) underway, the *Tbx5^+^/Mef2cAHF^+^* lineage cells form a ring situated between the RV and LV primordia, which is observed from multiple perspectives including anterior, short-axis (SAx) and long-axis (LAx) views. **o-q**, At E8.5 (approximately 9 somites), *Tbx5^+^/Mef2cAHF^+^* lineage cells are observed in a portion of the AV canal primoridium (pAVC and arrowheads). **r-t**, At E10.5, a band of *Tbx5^+^/Mef2cAHF^+^* lineage cells extends from the interventricular groove at the outer curvature to the inner curvature of the heart. SAx and LAx show that *Tbx5^+^/Mef2cAHF^+^* lineage cells occupies a crossroads for heart morphogenesis, spanning the growing interventricular septum to the AV canal (AVC), adjacent to the OFT. All scale bars equal 100 microns.

We further examined the lineage contributions of *Tbx5^+^/Mef2cAHF*^+^ septal progenitors at subsequent timepoints during cardiac morphogenesis. At the linear heart tube stage at E8.25, the labeled *Tbx5^+^/Mef2cAHF*^+^ lineage appeared intricately arranged into a ring of cells between the future left and right ventricles (Fig. 2j, k). This circlet configuration was maintained during rightward looping of the heart at E8.5 (Fig. 2o-q) and subsequent chamber formation. At E10.5, we observed a band of lineage-labeled cells from the IV groove at the outer curvature to the inner curvature near the AV groove that extended posteriorly to the atria (Fig. 2r-t), reminiscent of the “primary ring” ^9–11^ and consistent with oriented cell growth of the developing IVS ^12^. Optical sections showed the labeled lineage superiorly at a crossroads of the AV canal, outflow tract and atria (Fig. 2s). This populates a morphogenetic nexus, where VSDs can occur due to abnormal connections between the IVS and the AV canal, outflow tract cushions, or the muscular septum itself. In the atria, we noted lineage-labeled cells at the midline, superiorly and inferiorly (Supplemental Figure 1a-e), consistent with clonal growth in the body of the atria ^12^. This suggests that the *Tbx5^+^/Mef2cAHF^+^* lineage potentially marks the presumptive location of the IAS for later stages. Consequently, this data is consistent with a notion that the *Tbx5^+^/Mef2cAHF*^+^ lineage is configured early in heart development, well before subsequent morphogenetic events of chamber formation and cardiac septation.

### IVS disorganization and ventricular hypoplasia from cell ablation of *Tbx5*^+^/*Mef2cAHF*^+^ progenitors

To determine a role for the *Tbx5^+^/Mef2cAHF*^+^ septal progenitors during heart development, we conditionally ablated these cells at E6.5. To this end, we generated an intersectional recombinase-responsive *DTA176* knock-in mouse allele at the *Hip11* safe-harbor locus, by which cells would be killed where approximately 100-200 molecules of DTA176 were expressed ^36,37^. Upon a single dose of tamoxifen administration to pregnant dams at E6.5, intersectional-*DTA* mutant (*Tbx5^CreERT2/+^; Mef2cAHF-DreERT2; Hip11^Intersectional-DTA176/+^*) embryos displayed RV hypoplasia (i.e. a reduction in RV chamber size) at E9.5 (Fig. 3a, b) and E12.5 (Fig. 3c-j), together with IVS disorganization and non-compaction with a blunted IV groove (Fig. 3h, j). Further, there were defects of the AV complex region and absence of the IAS (Fig. 3h, j). A few *Tbx5^+^/Mef2cAHF^+^* lineage cells remained in mutant embryos, likely reflecting some degree of inefficient dual recombination of both the reporter allele ^38,39^ and the intersectional-*DTA* transgene ^40^. Beyond E12.5, intersectional-*DTA* mutant embryos were not recovered. Results of *Tbx5*^+^/*Mef2cAHF*^+^ progenitor ablation contrasted with findings after the ablation of the LV-enriched *Hand1*^+^ lineage, which caused LV hypoplasia at E10.5 and full recovery of the LV by E16.5 ^41^. Hence, our findings suggested that the *Tbx5^+^/Mef2cAHF^+^* septal progenitors were not only important for IVS development and AV septation, but also for RV chamber formation.

**Figure 3.**
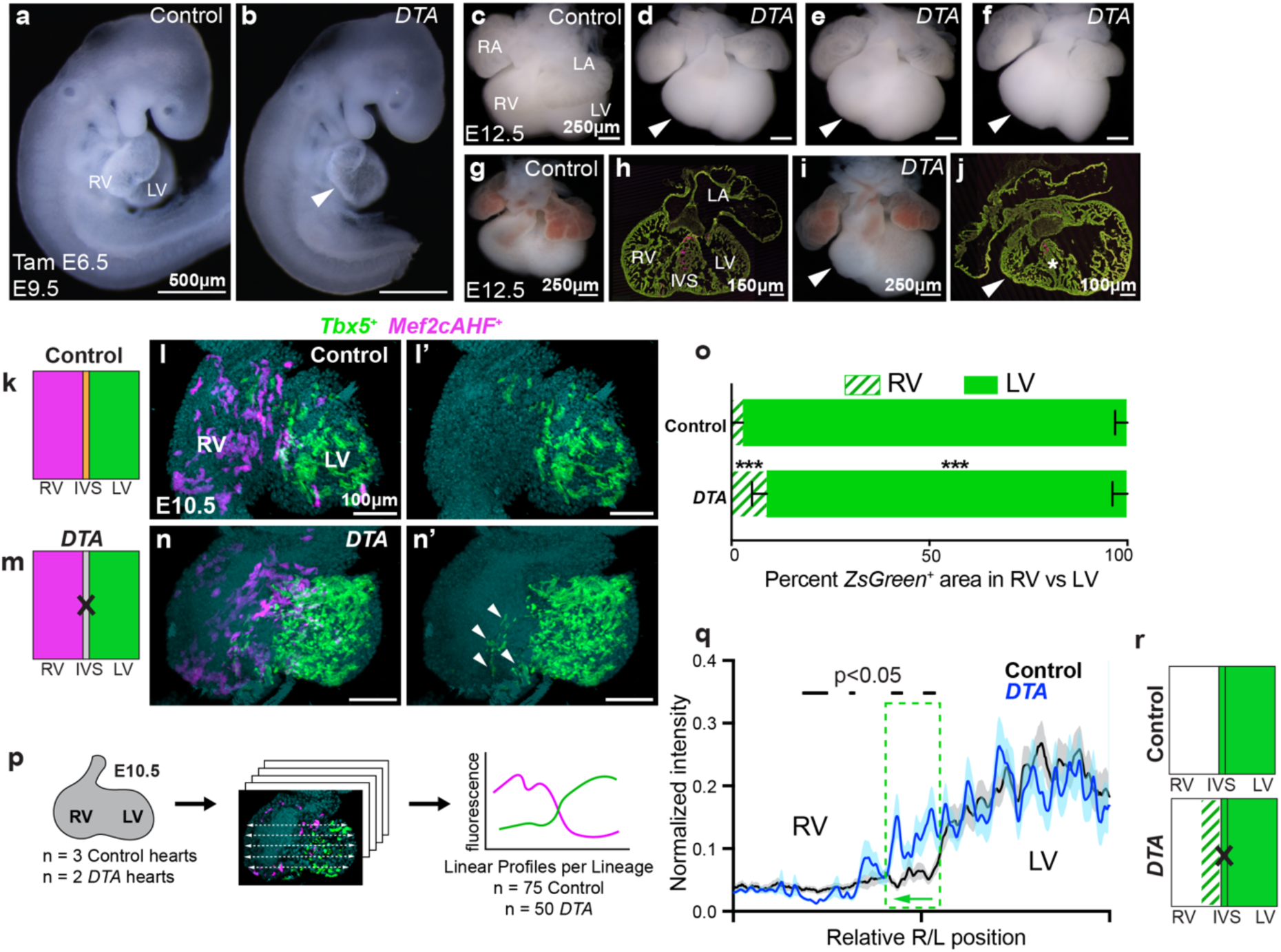
Cell ablation of *Tbx5^+^/Mef2cAHF*^+^ progenitors causes RV hypoplasia, IVS disorganization, and lineage mixing. **a-j**, Misexpression of *diphtheria toxin* (*DTA*) in *Tbx5^+^/Mef2cAHF*^+^ progenitors in *Tbx5^CreERT2^*^/+^;*Mef2cAHF-DreERT2*;*ROSA26^Ai66^*^/*+*^*;Hip11^intersectional-DTA/+^* embryos (*DTA*), compared to *Tbx5^CreERT2^*^/+^;*Mef2cAHF-DreERT2*;*ROSA26^Ai6^*^6/+^*;Hip11^+/+^* (control), resulted in right ventricle (RV) hypoplasia (arrows) at E9.5 (a, b; scale bars: 500 microns), and E12.5 (c-j; scale bars 250 microns), along with disorganization and non-compaction of the interventricular septum (IVS) (asterisk) (g-j). right atrium (RA), left atrium (LA). **k-n’**, At E10.5, mixing of *Tbx5*^+^ lineage (white arrows in n’ show ectopic *Tbx5*^+^ lineage ZsGreen cell in the right heart) was observed by lightsheet microscopy in *Tbx5^CreERT2^*^/+^;*Mef2cAHF-DreERT2*;*ROSA26^Ai6/Ai66b^;Hip11^intersectional-DTA/+^* embryos (*DTA*) (n=2) compared to *Tbx5^CreERT2^*^/+^;*Mef2cAHF-DreERT2*;*ROSA26^Ai6/Ai66b^;Hip11^+/+^* embryos (control) (n=3). (k, m) Cartoon depictions of experiment. **o**, Distribution of the *Tbx5*^+^ lineage by ventricular chamber demonstrates that the *Tbx5*^+^ lineage is increased in the RV of *DTA* mutants (p=0.000513 by two-sided T test), compared to controls. Cells of the RV and left ventricle (LV) were evaluated in five optical slices at different anterior-posterior planes per embryonic heart sample of 3 controls and 2 mutants. Mean and standard deviation are shown. **p-r**, Linear profiles quantified fluorescence intensity across the heart at the right or left (R/L) positions. *Tbx5*^+^ lineage cells expanded rightward into the RV (depiction in (r) by hashed area, dashed green box in (q) shows significant difference between traces, indicative of a rightward expansion (green arrow)), consistent with a compartment boundary disruption. Linear profiles at 5 apical-basal planes per optical slice, for 5 optical slices at different anterior-posterior planes per embryonic heart sample of 3 controls and 2 *DTA* mutants were assessed, as depicted in (p). Statistical significance (p<0.05) between linear profiles of control and *DTA* mutants was determined by Welch’s two-sided t-test at each position along the right-left axis. Mean and standard error of the mean are shown. Precise p-values can be found in the Source data.

### Lineage mixing from disruption of an IVS compartment boundary upon ablation of *Tbx5^+^/Mef2cAHF*^+^ septal progenitors

We hypothesized that the *Tbx5^+^/Mef2cAHF^+^* septal progenitors establish a compartment boundary at the IVS. Upon ablation of *Tbx5^+^/Mef2cAHF^+^* septal progenitors, we predicted that if either lineage expanded into the other ventricular chamber, then this would suggest lineage mixing from loss of compartment boundary integrity. To determine if *Tbx5^+^/Mef2cAHF*^+^ septal progenitors were essential for a compartment boundary at the IVS, we tested the effects of ablating *Tbx5^+^/Mef2cAHF*^+^ septal progenitors on lineage segregation (Fig. 3k-n’). We quantified the distribution of the *Mef2cAHF*^+^ or *Tbx5*^+^ lineages at E10.5 after a single dose of tamoxifen administration at E6.5 using two metrics. First, we assessed each fluorescently-labeled lineage by distribution in the RV or LV. We observed that cells of the *Mef2cAHF*^+^ lineage were enriched in the RV of control embryos (*Tbx5^CreERT2/+^;Mef2cAHF-DreERT2;Hip11^+/+^*), and this distribution was not different in mutant intersectional-*DTA* embryos (*Tbx5^CreERT2/+^;Mef2cAHF-DreERT2;Hip11^Intersectional-D^*^TA176^*^/+^*) (Extended Data Fig. 2a-c). Cells of the *Tbx5*^+^ lineage were normally enriched in the LV and nearly absent in the RV of controls. However, in mutant intersectional-*DTA* embryos, a larger percentage of *Tbx5*^+^ lineage cells were located in the RV (Fig 3o).

Second, we quantified the fluorescence intensity as a linear profile across the ventricular chambers by lightsheet microscopy (Fig. 3p). Cells of the *Mef2cAHF*^+^ lineage were again enriched in the RV in controls, and showed reduced fluorescence in the LV in mutant intersectional-*DTA* embryos (Extended Data Fig. 2d-f). Moreover, cells of the *Tbx5^+^* lineage were again highly enriched in the LV and rarely found in the RV. In contrast, cells of the *Tbx5*^+^ lineage in intersectional-*DTA* mutants were observed more rightward, including in the RV (Fig 3q, r). Therefore, we inferred that *Tbx5^+^/Mef2cAHF*^+^ progenitors were essential for preventing lineage mixing between the RV and LV, by maintaining compartment boundary integrity at the developing IVS. This data may imply preferential regulation of first heart field derivatives over the adjacent second heart field derivatives.

### Heterozygous loss of *Tbx5* in the IVS leads to VSDs

*TBX5* is a transcription factor gene that causes VSDs in Holt-Oram Syndrome from haploinsufficiency in humans ^42–44^ and mice ^45^. We wondered if heterozygous loss of *Tbx5* in the IVS would cause VSDs. We used the *Mef2cAHF-Cre* ^46^ in combination with a conditional deletion of *Tbx5* (*Tbx5^flox^*) ^45^, to conditionally delete *Tbx5* heterozygously in a domain that overlaps with *Tbx5* expression at the IVS ^13^. We observed membranous VSDs in *Mef2cAHF-Cre*;*ROSA26^mTmG/+^;Tbx5^flox/+^* mutant embryos (n=4/7) compared to controls (n=0/3), at E14.5 (Extended Data Fig. 3a-c). This provided evidence that appropriate *Tbx5* dosage in the IVS was essential for proper ventricular septation. Whether *Tbx5* is required in *Mef2cAHF*^+^ progenitors or in a subsequent stage for IVS formation remains to be determined.

### Disturbed septal lineage contributions from reduced *Tbx5*

As TBX5 dosage reduction globally or only in the IVS results in VSDs, we reasoned that reducing TBX5 dosage may affect the regulation of the *Tbx5^+^/Mef2cAHF^+^* septal progenitors and their subsequent lineage contributions. We evaluated a reduction of *Tbx5* using a hypomorphic (*Tbx5^CreERT2^*) allele ^44^, which has been used thus far in this study to follow the *Tbx5*^+^ and *Tbx5*^+^*/Mef2cAHF*^+^ lineages, in combination with a conditional deletion of *Tbx5* (*Tbx5^flox^*) ^45^ at E6.5. This resulted in levels of *Tbx5* that are estimated to be about 25% of wildtype in the *Tbx5^+^* lineage of *Tbx5* mutants. In control embryos (*Tbx5^CreERT2/+^;Mef2cAHF-DreERT2;ROSA26^Ai66/Ai6^*), the *Tbx5^+^/Mef2cAHF*^+^ septal lineage overlapped at the lineage front of the *Tbx5*^+^ lineage in the IVS, from the base to the apex of the heart at the IV groove, spanning from anterior to posterior of the heart in the IVS (Figure 4a-a”, Extended Data Fig. 4a, b, Video 2). Among *Tbx5^CreERT2/flox^* mutant embryos (*Tbx5^CreERT2/flox^;Mef2cAHF-DreERT2;ROSA26^Ai66/Ai6^*), we found that reduced TBX5 dosage at E6.5 caused a spectrum of defects, frequently including VSDs, ASDs or AV canal defects at E14.5 (Fig. 4b-d’), reminiscent of features of Holt-Oram Syndrome ^42,43,47^. The normally organized distribution of the tdTomato^+^ cells was highly irregular in the *Tbx5^CreERT2/flox^* mutant hearts regardless of VSDs (Extended Data Fig. 4a-h). tdTomato^+^ cells were less apparent in the posterior IVS (Fig. 4a’, a”, b’, b”, c’, c”, d’) and IAS (Supplemental Figure 2), and tdTomato^+^ cells were nearly absent in a heart that displayed a severe AV canal defect and chamber hypoplasia (Fig. 4d, d’, Extended Data Fig. 4g, h).

**Figure 4.**
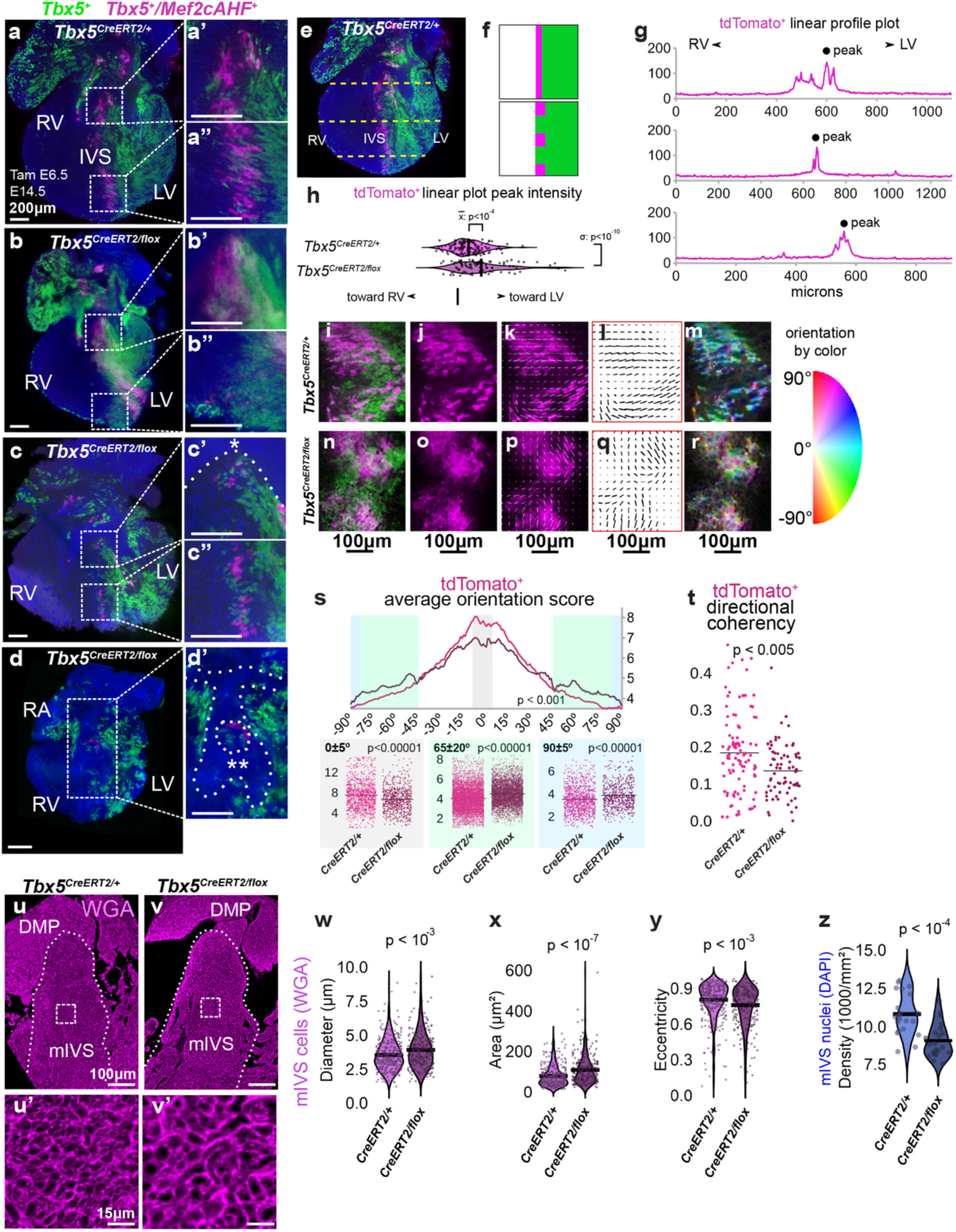
Reduced TBX5 dosage caused ventricular septal defects (VSDs), perturbations to IVS boundary position and integrity, and abnormal IVS cell arrangement. **a,** Mid-posterior optical sections from lightsheet microscopy of a control (*Tbx5^CreERT2/+^*;*Mef2cAHF-DreERT2*;*ROSA26^Ai6/Ai66^*) heart display *Tbx5^+^* lineage (ZsGreen) and *Tbx5^+^/Mef2cAHF^+^* lineage (tdTomato immunostaining) cells. *Tbx5^+^/Mef2cAHF^+^* lineage cells in the interventricular septum (IVS) extended from the base (a’) to the apex (a”) of the heart, where cells were highly organized. (B-D) A spectrum of phenotypes of *Tbx5^CreERT2/flox^* mutants (*Tbx5^CreERT2/flox^;Mef2cAHF-DreERT2*;*ROSA26^Ai6/Ai66^*) were observed, including intact IVS (b-b”), membranous VSD (c-c”, asterisk) and atrioventricular septal defect (double asterisk in d’). **b’, b”, c’, c”, d’**, A maldistribution and disorientation of *Tbx5^+^/Mef2cAHF^+^* lineage cells were observed in *Tbx5^CreERT2/flox^* mutants. (a-d’) sale bars: 200um. **e-h**, Linear plot profiles at positions along the anterior-posterior and apical-middle-basal axis (dashed lines) of control (116 profiles from 4 samples) and *Tbx5^CreERT2/flox^* mutant (87 profiles from 3 samples) hearts for *Tbx5^+^/Mef2cAHF^+^* lineage cells showed a leftward shift of boundary positioning and broadening of the *Tbx5^+^/Mef2cAHF^+^* lineage position, consistent with lineage mixing. The image in (e) is a repeat of the image in (a). **f**, Cartoon depiction. **g**, Examples of linear profile plots are shown, and (**h**) aggregated data is depicted and quantified. Statistics determined by two-sided unpaired F-test to compare variance (p=2.621e-11) and two-sided unpaired t-test (p=9.741e-05) to compare means. **i-s**, Orientation scores for cells from each channel (tdTomato^+^ or ZsGreen^+^) was delineated for each genotype, and distributions were plotted as a function of angle. *Tbx5^CreERT2/flox^* mutant hearts scored worse for orientation of *Tbx5^+^/Mef2cAHF^+^* lineage (tdTomato^+^) cells in the dominant direction (range from −5 to 5 degrees) and scored higher in the orientation orthogonal to the dominant direction (range from 85 to 95 degrees and −85 to −95 degrees), as determined in (s) by two-sided Watson U2 test across all orientations (p<0.001) or by two-sided Wilcoxon rank sum test with continuity correction for selected orientations (p=3.207e-10, 2.2e-16, 7.839e-09, for left-to-right plots). (i-r) scale bars: 100 microns. **t**, *Tbx5^+^/Mef2cAHF^+^* lineage (tdTomato^+^) cells demonstrated worse directional coherency scoring in *Tbx5^CreERT2/flox^* mutants (99 regions from 4 controls, 74 regions from 3 mutants) by two-sided Wilcoxon rank sum test with continuity correction (p=0.003845). **u-y**, Cell morphometry at E14.5 by wheat germ agglutinin (WGA) staining of cell membranes in the muscular IVS (mIVS) (dashed outline) of controls (3 planes per sample, 2 samples) and *Tbx5* mutants (3 planes per sample, 2 samples). scale bars: 100 microns (u, v); 15 microns (u’, v’). Images of (u-v’) are repeated in Supplemental Data Figure 3b, b’, f, f’. Quantification of cell (**w**) diameter, (**x**) area, and (**y**) eccentricity. **z**, Nuclei density by DAPI staining of the IVS. Statistics determined by two-sided unpaired T-tests (W: p=0.0002955, X: p=2.9e-08, Y: p=0.0007001, Z: p=5.296e-05).

We used quantitative morphometry to assess the position and distribution of the *Tbx5^+^/Mef2cAHF^+^* septal lineage at the IVS. We tested this by quantifying linear profiles of the labeled *Tbx5^+^/Mef2cAHF*^+^ septal lineage in the heart. We found that both distribution and position of the *Tbx5^+^/Mef2cAHF*^+^ septal lineage was disturbed in *Tbx5^CreERT2/flox^* mutants (Fig. 4e-h, Extended Data Fig. 4i-k), including an increase of tdTomato^+^ cells leftward in the LV, suggesting abnormal lineage position. As well, we observed a broader band of *Tbx5^+^/Mef2cAHF*^+^ septal lineage cells (Fig. 4g, h), suggesting that proper TBX5 dosage maintains integrity of the compartment boundary marked by the *Tbx5^+^/Mef2cAHF*^+^ lineage at the IVS. We also observed a reduction, or sometimes absence, of the labeled septal lineage among AV complex region cells, as well as remnant atrial septal tissue in *Tbx5^CreERT2/flox^* mutants (Fig. 4d, d’).

We further observed that cells of the *Tbx5^+^/Mef2cAHF*^+^ lineage in the IVS were arranged like a stack of coins, especially from the apex to the mid-septum (Fig. 4a”). We wondered if proper TBX5 dosage might be necessary for maintaining appropriate cell arrangement in the IVS. Therefore, cell alignment was determined and quantified by two metrics, average orientation score or directional coherency. In *Tbx5^CreERT2/flox^* mutant hearts, *Tbx5^+^/Mef2cAHF^+^* lineage (tdTomato^+^) and *Tbx5*^+^ lineage (ZsGreen^+^) cells were reduced in the dominant direction of cell orientation and were more frequently in the orientation orthogonal to the dominant direction (Fig. 4i-t, Extended Data Fig. 4l-s). Further, both lineages demonstrated worse directional coherency in *Tbx5^CreERT2/flox^* mutants (Fig. 4t, Extended Data Fig. 4u). In a complementary analysis, we also evaluated cell geometry and tissue architecture of the IVS using staining for cell borders by wheat germ agglutinin (WGA) and nuclei by DAPI. We found in IVS cardiomyocytes of *Tbx5* mutants that cell diameter and area was enlarged (Fig. 4u-x, Supplemental Figure 3a-h’), while cell eccentricity (a measure of a cell’s elongation) and nuclei density was reduced (Fig. 4y, z). Taken together, this evidence supports a notion that TBX5 is important for proper cell arrangement in the IVS.

### *Tbx5*-sensitive genes encode guidance cues

To find downstream effectors of Tbx5 that may mediate the regulation of *Tbx5^+^/Mef2cAHF*^+^ lineage cells, we applied scRNA-seq at E13.5, prior to completion of ventricular septation. We micro-dissected the RV, LV and IVS+AV complex regions in controls and *Tbx5* mutants (*Tbx5^CreERT2/flox^;Mef2cAHF-DreERT2;ROSA26^Ai66/Ai6^*) (Fig. 5a). These samples were labeled at E6.5 for progenitors of the *Tbx5^+^/Mef2cAHF*^+^ lineage (*tdTomato*^+^) and *Tbx5*^+^ lineage (*ZsGreen*^+^). In samples from each cardiac tissue region, we detected clusters enriched for *Tnnt2*^+^ cardiomyocytes (CMs), *Postn*^+^ fibroblasts, *Plvap*^+^ endothelial cells, *Tbx18^+^/Wt1^+^* epicardial cells, *Hba-x*^+^ red blood cells, and *C1qb*^+^ white blood cells (Fig. 5b, Extended Data Fig. 5A-i).

**Figure 5.**
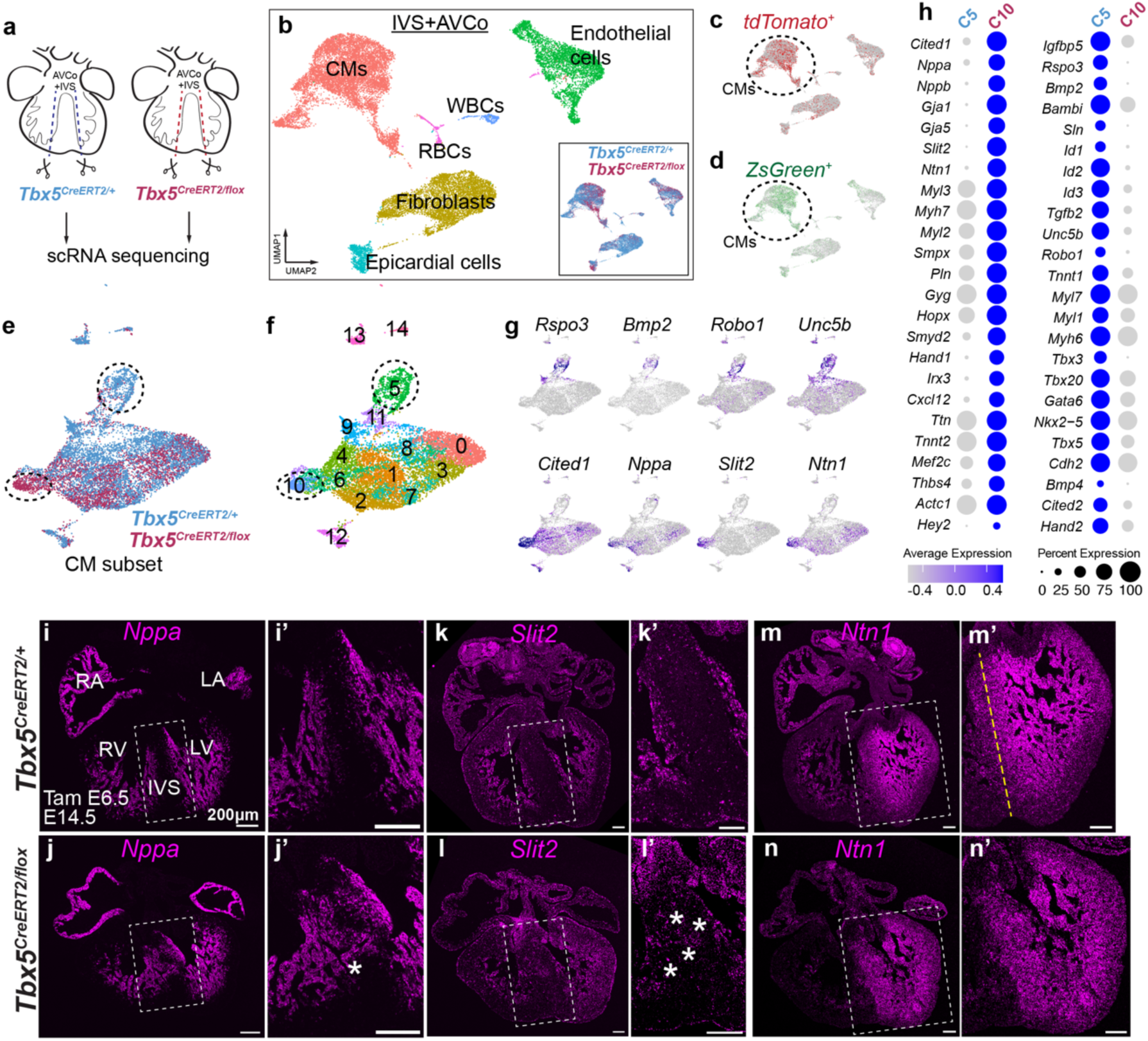
*Slit2* and *Ntn1* are *Tbx5*-sensitive genes in the interventricular septum (IVS). **a**, At E13.5, we micro-dissected the right ventricle (RV), left ventricle (LV) and IVS+atrioventricular complex (AVCo) regions in *Tbx5^CreERT2/+^* controls and *Tbx5^CreERT/flox^* mutants (*Tbx5^CreERT2/flox^;Mef2cAHF-DreERT2*;*ROSA26^Ai6/^*^Ai66^) with the labeled *Tbx5^+^/Mef2cAHF^+^* lineage (*tdTomato^+^*) and *Tbx5*^+^ lineage (*ZsGreen*^+^) cells, after a single dose of tamoxifen at E6.5. **b**, UMAP visualization of IVS+AVCo by cell type clusters and *Tbx5* genotype (inset) (controls, n=4; *Tbx5* mutants, n=2). CMs, cardiomyocytes; WBCs, white blood cells; RBCs, red blood cells. **c**, *ZsGreen^+^* and (**d**) *tdTomato*^+^ cells are enriched among CMs. **e**, UMAP shows a *Tnnt2^+^* CM subset by *Tbx5* genotype or (**f**) Louvain clusters. **g**, Feature plots show that AVCo region genes (*Rspo3, Bmp2, Robo1, Unc5b*) are enriched among control-enriched cluster 5, suggesting downregulation of these genes in *Tbx5* mutants. Trabecular genes (*Cited1, Nppa, Slit2, Ntn1*) were enriched in the *Tbx5* mutant-enriched cluster 10. **h**, Dot plots of selected genes that are upregulated in cluster 10 (C10) or in cluster 5 (C5). Significance of adj p<0.05 was determined by two-sided Wilcoxan rank sum. **i-n**, Fluorescent *in situ* hybridization of trabecular genes *Nppa*, *Slit2* and *Ntn1* in *Tbx5^CreERT2/+^* controls or *Tbx5^CreERT2/flox^* mutants at E14.5. **i, i’, k, k’**, *Nppa* and *Slit2* are normally expressed in the ventricular trabecular layer and excluded from the core of the IVS. **j, j’, l, l’**, In a *Tbx5^CreERT2/flox^* mutant with an intact IVS, *Nppa* and *Slit2* are misexpressed in the IVS (asterisks). **m, m’**, *Ntn1* is normally expressed in the trabecular layer and in a gradient across the IVS from left to right, demarcated by yellow dashed line. **n, n’**, In a *Tbx5^CreERT2/flox^* mutant with an intact IVS, *Ntn1* is expanded across the IVS, flattening its gradient. All scale bars equal 200 microns.

In particular, *tdTomato^+^* cells were most abundant among clusters of *Tnnt2*^+^ cardiomyocytes (CMs) of the IVS+AV complex regions (Fig. 5c). However, we were unable to identify a gene expression signature that was unique to the *Tbx5*^+^/*Mef2cAHF*^+^ lineage, beyond the lineage marker itself, at this stage. Likewise, *ZsGreen*^+^ cells were also enriched among clusters of *Tnnt2^+^* CMs in this region (Fig. 5d), indicative of the *Tbx5*^+^ lineage contribution to CMs in the region, as well as a site of conditional reduction of *Tbx5* among *Tbx5* mutants.

We then focused our analysis on a subset of *Tnnt2*^+^ enriched clusters from IVS+AV complex regions. We identified genotype-enriched clusters that were abundant with cells from controls (Cluster 5) or *Tbx5* mutants (Cluster 10) (Fig. 5e, f). In the mutant-enriched cluster 10, genes typically expressed in ventricular trabeculae were enriched (Fig. 5g, h). These included natriuretic peptide *Nppa*, transcription factor *Cited1,* and bone morphogenetic protein *Bmp10*, suggesting ectopic expression of these trabecular genes in the IVS of *Tbx5* mutants. Intriguingly, guidance cues *Netrin1* (*Ntn1)* and *Slit2,* which are best known for axonal development or vasculogenesis ^48^, were also dysregulated (Fig. 5g, h, Extended Data Fig. 5j,k, Supplemenatal Figure 4a, b). Notably, *SLIT2* is implicated as a CHD-risk gene in mice and humans ^49–52^. For *NTN1*, it’s link to human CHD requires further clarification, as a copy number variation by intragenic deletion of *NTN1*, along with heterozygous loss of 22q11.21 and gain of the Y chromosome, was associated with septal defects in a patient ^53^. Control-enriched cluster 5 included genes expressed in the AV complex region, encoding the Wnt co-receptor *Rspo3*, morphogen *Bmp2*, and transcription factors *Tbx2* and *Tbx3*, and *Cdh2*, which encodes N-Cadherin, suggesting that these genes are reduced in *Tbx5* mutants (Fig. 5g, h). As well, the SLIT-receptor *Robo1,* which is implicated in mouse and human VSDs ^49,51,54^, and Netrin-receptor *Unc5b,* were also downregulated in *Tbx5* mutant cells (Fig. 5g, h).

We examined differential gene expression between controls or *Tbx5* mutants in *tdTomato*^+^ (*Tbx5^+^/Mef2cAHF^+^* septal lineage) or *ZsGreen*^+^ (*Tbx5^+^* lineage) CMs. We found many differentially expressed genes that overlapped among comparisons of *tdTomato*^+^ or *ZsGreen*^+^ clusters, including *Nppa*, *Cited1,* and *Slit2*, consistent with findings from cluster-to-cluster comparisons (Extended Data Fig. 5j, k).

We used orthogonal assays to validate candidate TBX5-sensitive genes. By fluorescence *in situ* hybridization in formalin fixed paraffin embedded (FFPE) sections, we observed a reduction of gene expression of *Bmp2, Robo1 and Unc5b* in the AV complex region of mutant hearts when compared to controls. (Extended Data Fig. 6a-f”). We found that the IVS-enriched gene *Irx2* was reduced in the IVS of *Tbx5* mutants (Extended Data Fig 6g-h’). As well, we detected expansion of *Nppa* and *Slit2* into the IVS and compact layer of *Tbx5* mutant hearts, both of which are normally expressed only in the trabecular regions of both ventricles of control hearts (Fig. 5i-l’). Moreover, we discovered *Ntn1* expression enriched in LV trabecular CMs and in a gradient across the control IVS (Fig. 5m, m’, Extended Data Fig. 6m-m”), consistent with immunostaining of NTN1 in the heart at E14.5 (Extended Data Fig. 6o). However, in *Tbx5* mutants, the *Ntn1* gradient was flattened and expanded in the IVS by reduced TBX5 dosage (Fig. 5n, n’). Interestingly, we observed abnormal *Nppa*, *Slit2* and *Ntn1* in *Tbx5* mutants with or without VSDs, suggesting that this dysregulated gene expression was not secondary to VSDs (Extended Data Fig. 6i-n”). Notably, earlier at E11.5, *Ntn1* was LV-enriched during normal ventricular septation and broadly expanded into the RV of *Tbx5* mutants, while *Slit2* appeared unchanged in *Tbx5* mutants (Extended Data Fig. 7a-h). In addition, analysis of ChIP-seq of TBX5 from embryonic mouse hearts ^55^ showed TBX5 occupancy at promoters of *Nppa, Slit2*, and *Ntn1* (Extended Data Fig. 8a-c), suggesting these guidance genes are likely to be direct targets of TBX5 in the heart.

### *Tbx5*-*Slit2* and *Tbx5*-*Ntn1* genetic interactions

We evaluated if genetic interactions existed between *Tbx5* and *Slit2* or *Ntn1* (Extended Data Fig. 9). As *Tbx5* heterozygous mutant (*Tbx5^del/+^)* ^45^ mouse embryos display muscular or membranous VSDs, we wondered if heterozygous *Slit2* or *Ntn1* loss of function (LOF) mutant embryos might alter *Tbx5*-dependent phenotypes. We mated *Tbx5^del/+^* mice with either *Slit2* (*Slit2^+/-^)* or *Ntn1 (Ntn1^beta-actin-Cre/+^* referred here subsequently as *Ntn1^+/-^* ^56,57^) heterozygous LOF mice. In *Tbx5^del/+^* embryos at E14.5, we observed membranous (n=5/8) or muscular (n=3/8) VSDs by histology (Extended Data Fig. 9a, b, e), consistent with previous reports ^45^. Among *Slit2^+/-^* embryos at E14.5, we observed a membranous VSD (n=1/8) (Extended Data Fig. 9c, e). This contrasted with a previous report that did not detect any VSDs in *Slit2^+/-^* albeit using a different mutant *Slit2* allele ^49^. In *Tbx5^del/+^*;*Slit2^+/-^* compound heterozygous embryos, we observed a non-significant effect on the reduction of membranous VSDs (n=5/8) (log OR −2.5; p=0.09 in a Generalized Linear Model without adjustments for multiple comparisons) (Extended Data Fig. 9d, e), relative to the expected incidence of membranous VSDs if there was no genetic interaction between *Tbx5* and *Slit2*.

We next tested if there was a genetic interaction between *Tbx5* and *Ntn1* (Extended Data Fig. 9f-j). In *Ntn1^+/-^*embryos, we observed a membranous (n=1/9) or a muscular (n=1/9) VSD. In *Tbx5^del/+^*;*Ntn1^+/-^* compound heterozygous embryos, we observed a significant decrease of membranous VSDs (n=1/8; log OR −5.3; p=1x10^-3^ by a Generalized Linear Model without adjustments for multiple comparisons), relative to the expected incidence of membranous VSDs if there was no genetic interaction between *Tbx5* and *Ntn1,* but not significant changes to prevalence of muscular VSDs (n=4/8; log OR 0.12; p=0.93), consistent with a genetic interaction between *Tbx5* and *Ntn1* for ventricular septation.

We wondered if there were other quantitative histologic differences, in addition to the changes in incidence of VSD types. To quantify morphometry from histology, we leveraged a deep learning algorithm, known a CODA ^58^, to detect embryonic mouse heart components from hematoxylin and eosin (H&E) sections. We generated three-dimensional (3-D) tissue reconstructions, to visualize and quantify anatomic structures and CHDs, including membranous and muscular VSDs, at cellular resolution (Fig. 6a-e). Using this machine learning method, we discovered that the cell density of the IVS, which we termed IVS fill, was increased in *Tbx5^del/+^*; *Ntn1^+/-^* embryos compared to wildtype (p<0.05 by Fisher’s exact test), while classification of the IVS as trabecular was significantly reduced in *Tbx5^del/+^*;*Ntn1^+/-^* embryos (p<0.05 by Fisher’s exact test) (Extended Data Fig. 9m). In addition, we estimated the minimal area for each membranous or muscular VSD, when present (Extended Data Fig. 9o, p). In *Tbx5^del^*^/+^;*Slit2^+/-^* compound mutants, we detected statistically significant levocardia (p<0.001 by Fisher’s exact test), as determined by the axis of the heart compared to the spine and sternum (Extended Data Fig. 9q). This uncovered a genetic interaction between *Tbx5* and *Slit2* related to heart position, demonstrating that machine learning-based morphometry can quantify unexpected anatomic findings.

**Figure 6.**
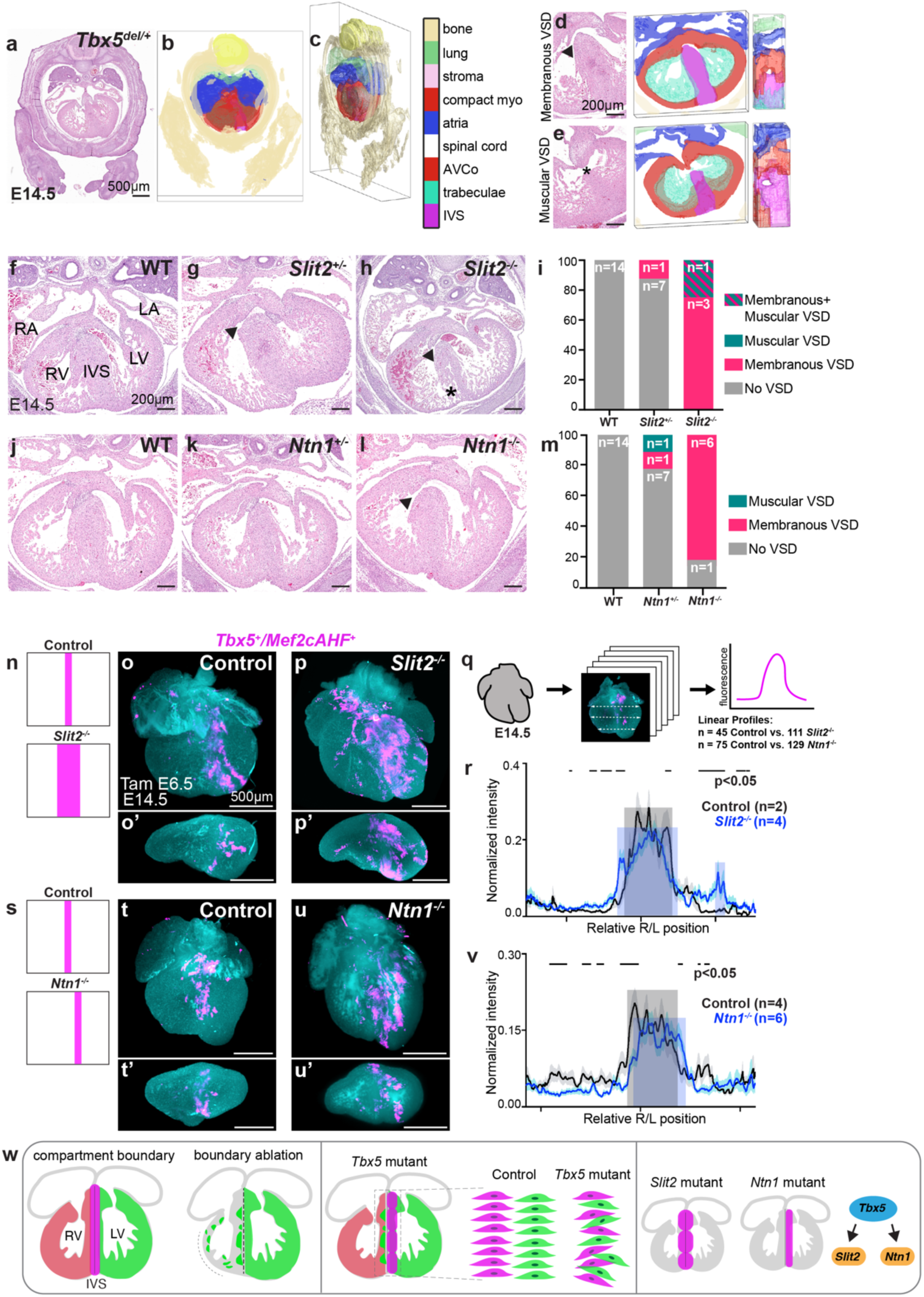
*Slit2* and *Ntn1* are essential for proper ventricular septation and compartment boundary regulation. **a-e**, Machine learning-based 3-D reconstructions of the embryonic heart and other tissues from histology at E14.5 were used for quantitative morphometric analysis of the embryonic heart (Extended Data Fig. 9, Extended Data Fig. 10). Compact myo, compact myocardium; AVCo, atrioventricular complex region; IVS, interventricular septum. **d, e**, A 3-D reconstruction of a membranous ventricular septal defect (VSD) anteriorly and muscular VSD posteriorly from a *Tbx5^del^*^/+^ mutant heart. **f-h**, Histology and (**i**) incidence of muscular or membranous (arrowhead) ventricular septal defects (VSDs) of *Slit2* mutants are shown. Membranous VSDs were enriched in *Slit2^-/-^* (n=3/4, log OR 5.0, p=0.02 by a Generalized Linear Model without adjustments for multiple comparisons) and *Slit2^+/-^* (n=1/8, OR 2.5, p=0.02). Asterisk demarcates non-compacted interventricular septum (IVS). **j-l**, Histology and (**m**) incidence of muscular or membranous ventricular septal defects (VSDs) of *Ntn1* mutants are shown. Membranous VSDs were enriched in in *Ntn1^-/-^* (n=6/7, log OR 6.32, p=0.0052 by a Generalized Linear Model without adjustments for multiple comparisons) and *Ntn1*^+/-^ (n=1/9, OR 3.17, p=0.0052 by a Generalized Linear Model without adjustments for multiple comparisons). **n-r**, Lightsheet imaging of *Slit2*^-/-^ mutant (*Tbx5^CreERT2/+^*;*Mef2cAHF-DreERT2*;*ROSA26^Ai66/+^;Slit2*^-/-^)(n=4) hearts at E14.5 showed a broadened distribution of the *Tbx5^+^/Mef2cAHF*^+^ lineage (tdTomato^+^ immunostaining) compared to controls (*Tbx5^CreERT2/+^*;*Mef2cAHF-DreERT2*;*ROSA26^Ai66/+^;Slit2*^+/+^)(n=2), as (**r**) quantified by linear profiles of fluorescence signals. Gray shading represents control signal, and periwinkle represents mutant signal. Significance of p<0.05 was determined by Welch’s two-sided t-test at each position along the right-left axis. **s-v**, Lightsheet imaging of *Ntn1*^-/-^ mutant (*Tbx5^CreERT2/+^*;*Mef2cAHF-DreERT2*;*ROSA26^Ai66/+^;Ntn1*^-/-^)(n=6) hearts at E14.5 showed a leftward shift in positioning of the *Tbx5^+^/Mef2cAHF*^+^ lineage compared to controls (*Tbx5^CreERT2/+^*;*Mef2cAHF-DreERT2*;*ROSA26^Ai66/+^;Ntn1*^+/+^)(n=4), as (**v**) quantified by linear profiles. Significance of p<0.05 was determined by Welch’s two-sided t-test at each position along the right-left axis. Precise p-values are available in Source data. **n, s**, Cartoon depictions. (o’, p’, t’, u’) are optical sections that are orthogonal views to (o, p, t, u). **q**, For each sample, three linear profiles (basal, middle, apical) spanned the ventricular chambers from right to left from optical sections every 50 um from anterior to posterior of the heart. Precise values of statistics for (r) and (v) can be found in Source data. **w**, Cartoon depicts summary of findings. All scale bars are 200 microns.

### *Slit2* and *Ntn1* for ventricular septation

To determine if *Slit2* or *Ntn1* are essential for ventricular septation, we evaluated homozygous LOF mouse mutants for *Slit2* or *Ntn1* for VSDs. In *Slit2^-/-^* embryos at E14.5, we observed that homozygous loss of *Slit2* displayed complete penetrance for VSDs, including membranous (n=3/4; log OR 5.0, p=0.02 by a Generalized Linear Model) or muscular VSD (n=1/4) (Fig. 6f-i). We detected a statistically significant non-compacted IVS, which was determined by reduced IVS fill, and thinning of the right ventricular compact layer, and we quantified the area of membranous VSDs (Extended Data Fig. 10e-k). In addition, *Slit2* heterozygous LOF displayed an increase of membranous VSDs (n=1/8, OR 2.5, p=0.02 by a Generalized Linear Model), relative to the expected incidence of membranous VSDs in controls. This demonstrated a far higher incidence of VSDs in *Slit2* mutants than reported using an alternative *Slit2* LOF allele ^49^.

*Ntn1* homozygous LOF embryos displayed an increase of membranous VSDs (n=6/7; log OR 6.32, p=0.0052 by a Generalized Linear Model without adjustments for multiple comparisons) that we quantified by area, while other morphometry parameters were not significant (Extended Data Fig. 10f-r). *Ntn1* heterozgyous LOF embryos were enriched for membranous VSDs (n=1/9, OR 3.17, p=0.0052 by a Generalized Linear Model), relative to the expected incidence of membranous VSDs in controls. Collectively, this morphologic evidence implicates *Slit2* and *Ntn1* in proper ventricular septation during heart development.

As *Slit2* and *Ntn1* were necessary for ventricular septation, we asked if loss of *Slit2* or *Ntn1* might disturb distribution of the *Tbx5^+^/Mef2cAHF^+^* lineage in the IVS. In homozygous *Slit2* LOF mutants with the lineage reporter (*Tbx5^CreERT2/+^*;*Mef2cAHF-DreERT2*;*ROSA26^Ai^*^66^*^/+^*;*Slit2^-/-^)*, we observed a statistically-significant broadened distribution of the *Tbx5^+^/Mef2cAHF^+^* lineage compared to controls (Fig. 6n-r), consistent with lineage mixing from boundary disruption and implicating a role for SLIT2 in maintaining boundary integrity. Conversely, homozygous *Ntn1* LOF mutants with the lineage reporter (*Tbx5^CreERT2/+^*;*Mef2cAHF-DreERT2*; *ROSA26^Ai66/+^*;*Ntn1^-/-^)* demonstrated leftward displacement of the *Tbx5^+^/Mef2cAHF^+^* lineage compared to controls (Fig. 6s-v), implying that NTN1 signaling precisely positions the compartment boundary at the IVS. Taken together, this evidence suggested that SLIT2- and NTN1-signaling, as part of a Tbx5-dependent pathway, regulates the position and integrity of the compartment boundary labeled by the *Tbx5^+^/Mef2cAHF^+^* lineage (Fig. 6w).

## Discussion

We show that *Tbx5*^+^/*Mef2cAHF*^+^ progenitors prefigure a compartment boundary that becomes located at the junction between the left and right sides of the IVS, providing a cellular framework during development for heart patterning. Our study underscores a fundamental principle that early developmental events preconfigure structure and function of the heart and are susceptible to genetic risks that cause CHDs. Further, our results demonstrate developmental regulation of a compartment boundary by a disease-associated TF and that aberrant tissue patterning from boundary disturbances may be an etiology of birth defects. This study reiterates the importance of compartment boundaries in tissue patterning, provides evidence for compartment boundary disruptions as an etiology of birth defects, and adds to the few examples of compartment boundaries that exist in mammals.

We surmise that the rarity of compartment boundaries in mammals likely stems from a few reasons. The discovery of a compartment boundary was made by studying the fate of marked mosaic cells and their clonal segregation in the wing disk of the fly embryo ^59^. Subsequent examples of compartment boundaries were identified by broad genetic screens in *Drosophila* ^60^. In mammals, compartment boundaries were discovered by lineage analysis informed by another organism ^33^ or serendipitously ^30^, as we have done here. Compartment boundaries have been defined by several criteria. First, a compartment boundary segregates juxtaposed cell populations such as distinct cell lineages to restrict cell intermingling, and ablation of the boundary leads to cell mixing. Second, the expression domain of a selector gene corresponds with a compartment domain, loss of the selector gene eliminates the identity in this territory, and ectopic expression of the selector induces this identity. Third, a selector gene establishes a signaling center to maintain the compartments and boundary. Our results satisfy all of these criteria: Ablation of the *Tbx5*^+^/*Mef2cAHF*^+^ lineage disrupts segregation of *Tbx5*^+^ lineage cells in the LV from the *Tbx5*^-^ lineage in the RV of the developing heart. Reduced *Tbx5* expression impairs the integrity of the boundary and its patterning, and in previous work misexpression of TBX5 eliminates IVS formation ^13^. Third, TBX5 patterns signaling molecules that are important for morphologies that impact the boundary. In this context, we postulate that *Tbx5* may function as a candidate selector gene. Further experiments will be needed to formally ascertain this.

In this context, we identify a *Tbx5*-dependent pathway that regulates *Slit2* and *Ntn1*. Normally, TBX5 represses their ectopic expression in the IVS and compact layer. Both SLIT2 and NTN1 are guidance or adhesive cues in several developmental contexts. Complete loss of *Slit2* causes perinatal lethality (Plump et al., 2002). *Ntn1^-/^*^-^ die around E11.5 ^61^ or E14.5 ^56^. Between E10.5 and E14.5, homozygous *Ntn1*^-/-^ embryos exhibited absence of ventral spinal commissure, and disorganized dorsal root entry zone ^56^.

Here, we provide evidence that *Slit2* and *Ntn1* are direct targets of TBX5, and that *Tbx5* and *Slit2* or *Ntn1* genetically interact. Further, we found that *Slit2* and *Ntn1* are essential for proper ventricular septation, as well as compartment boundary integrity and positioning, respectively. Like TBX5, proper ventricular septation may be sensitive to imbalanced dosage of SLIT2 or NTN1. Of note, mutations of *SLIT2* and the SLIT-receptor *ROBO1* are associated with CHDs in mouse and humans ^49–52^. For NTN1, knockdown of *ntn1a* in zebrafish results in cardiovascular abnormalities ^53^, while *NTN1* had not yet been clearly implicated as a risk gene for mouse ^61^ or human CHDs ^53^. Even so, *DSCAM*, which encodes a NETRIN co-receptor, is in the risk region for Trisomy 21 ^62^, which often includes atrioventricular septal defects. Moreover, single nucleotide variants in *NEO1* and *UNC5B*, which encode NTN-receptors, are associated with CHDs ^63,64^. As mouse embryos from *Unc5b* loss, formerly known as *Unc5h2*, died of vascular abnormalities of the placenta by E12.5 ^65,66^, it remains unclear what role *Unc5b* might have in cardiac chamber formation and ventricular septation. As *Ntn1* expression is enriched in the LV and left-side of the IVS in the *Tbx5*^+^ compartment, we propose a working hypothesis that NTN1 may function as a TBX5-dependent signaling center for compartment boundary regulation.

The spatiotemporal position of the boundary is at the crux of cardiac septation and, when disrupted, spans sites for congenital cardiac anomalies. The sensitivity of the *Tbx5*^+^/*Mef2cAHF*^+^ lineage-labeled boundary to genetic perturbations, specifically *Tbx5* deficiency, is of significance in the context of CHDs. Such deficiency selectively compromises the boundary’s integrity and position, along with disturbances to cell orientation, leading to VSDs or AVSDs. These observations suggest that patterning during heart morphogenesis is exceptionally vulnerable to disruptions of gene dosage in early cardiac precursors. Early susceptibility could explain why CHDs arise at higher frequency than other birth defects. Thus, the *Tbx5*^+^/*Mef2cAHF*^+^ lineage provides a potential “beacon” for understanding the cellular basis of septation defects. Whether additional genetic perturbations that cause VSDs or AVSDs disrupt the compartment boundary remains to be determined. Likewise, it will be interesting to evaluate if the *Tbx5*^+^/*Mef2cAHF*^+^ lineage-labeled boundary corresponds with abnormal shifts in IVS position, like those observed in the endothelial loss of *Hand2*, which displays rightward shifted IVS and double inlet left ventricle (DILV) or two IVS primordia ^67,68^.

Several limitations of this study should be considered. Using inducible genetic tracing, there is an inherent variability from recombination of the fluorescent reporters within samples. For some comparisons, more samples of each genotype could have mitigated this limitation, but obtaining additional samples have been technically challenging, in large part due to a requirement of 5 alleles to be present. Notwithstanding, we have tried to account for recombination efficiency, when possible, in imaging analyses. We have made great effort to mitigate this concern by not quantifying measurements that would be greatly skewed by recombination efficiency. For this reason, for example, we have refrained from commenting on whether a given lineage is quantitatively increased or decreased if fewer samples were available. Instead, we have focused our analyses on the location of the cells, as we feel that this is less affected by recombination efficiency. Using orthogonal approaches, we measure in an unbiased and statistically robust fashion using various metrics of quantitative morphometry from lightsheet or histology images. Our interpretations are based on statistical robustness from a large number of sampled regions per embryo, in the comparisons that are presented. The repeated measurements are akin to the (much larger) measurements in single cell RNAseq.

Broadly, our research demonstrates the delicate interplay of progenitor cell behavior, gene expression and compartment boundary formation that determines heart development and genetic susceptibility that increases the risk of CHDs. This evidence further underscores the significance of early developmental stages for the ontogeny of some CHDs. Whether the extent of interaction between progenitor fields, migration of heart progenitors, or formation of the boundary may contribute stochastically to phenotypic variation observed as a frequent feature of CHD genetics in mice or humans remains unclear. The implications of these discoveries likely extend beyond cardiac morphogenesis, as these findings offer insights for the roles of compartment boundaries during mammalian development and uncover a vulnerability for organ patterning from genetic disturbances.

## Methods

### Mouse Lines

All mouse protocols (AN203375-00H and AN199784-00E) were approved by the Institutional Animal Care and Use Committee at UCSF. Mice were housed in a barrier animal facility with standard husbandry conditions at the Gladstone Institutes. Mice of *Tbx5^CreERT2IRES2xFLAG^* (abbreviated here as *Tbx5^CreERT2^*) and *Mef2cAHF-DreERT2* ^25^, *Tbx5^del/+^* and *Tbx5^flox/+^* ^45^, *ROSA26^Ai^*^66^ and *ROSA26^Ai6^* ^35^ were described previously. *Mef2cAHF-Cre* mice ^46^ were obtained from Brian Black (University of California, San Francisco). *Slit2^+/-^* mice (MMRC, Strain 065588-UCD, donated by Kent Lloyd, UC Davis) were generated by CRISPR/Cas9-targeted constitutive deletion of exon 8 and flanking splicing regions of *Slit2*. *Ntn1^+/-^* mice were derived from matings of *Ntn1* floxed mice (*Ntn1^flox/+^*; Jackson Laboratory #028038) ^56^ to *beta-actin-Cre* ^57^, which were obtained from Gail Martin (University of California, San Francisco). All mouse strains were maintained in the C57BL6/J background (Jackson Laboratory #664), except for *Tbx5^del^*^/+^, which was maintained in Black Swiss (Charles River, Strain Code 492), and *Slit2^+/-^* which was maintained in C57BL6/N (Jackson Laboratory, #005304). Both male and female embryos were collected from timed matings and used at random for experiments. Tamoxifen (Sigma-Aldrich; T5648) was suspended in sesame oil at 10mg/ml. The injected dose was 100ug per 1 gram of body weight.

### Cloning and Generation of Mouse Lines

We generated an attenuated diptheria toxin (DTA176) transgenic knock-in mouse under the control of the dual-recombinase intersectional cassette. DTA176 encodes an attenuated form of toxic fragment A from diptheria toxin ^36,37^, which requires ∼100-200 molecules for cell killing. This strategy was an effort to reduce potential problems of leaky DTA expression prior to recombinase-mediated activation. Briefly, *DTA176* ^37^ was cloned downstream of ROX-STOP-ROX and LOX-STOP-LOX sites to create a Dre- and Cre-dependent expression construct, which was derived from the Dre- and Cre-dependent tdTomato expression cassette of the Ai66 (RCRL-tdT) targeting vector ^35^. After the tdTomato cassette was removed using MluI sites and replaced by DTA176, the construct was cloned into the TARGATT targeting construct pBT346.3 (Applied Stem Cells) to target the transcriptionally inactive *Hip11* locus ^69^ using PacI and SmaI restriction enzyme sites. The final construct was verified using restriction digestion and Sanger sequencing. DNA was purified and injected along with mRNA for the Phi31o transposase according to manufacturer’s protocol to generate intersectional-DTA mutant (*Hip11^Rox-STOP-Rox-Lox-STOP-Lox-DTA1^*^76^) mice. To generate Ai66b mice (*ROSA26^Ai66b^*), which is a Dre-dependent tdTomato reporter, the LOX-STOP-LOX sites were removed from the Ai66 (RCRL-tdT) targeting vector ^35^, and then the transgene cassette was inserted into endogenous genomic loci via homologous recombination, as previously described ^70^.

### Whole Mount Embryo and Heart Immunofluorescence Labeling

Embryos were dissected from timed pregnant dams, including removal of yolk sac for genotyping, as previously described ^25^. Embryos staged up to E10.5 were fixed for 4 hours in 4% PFA at room temperature, then incubated with gentle rocking in clearing solution (8% SDS in 0.2M sodium borate buffer, pH 8.5) at 37°C for 1-3 hours until clear. E13.5 to E14.5 hearts were micro-dissected from freshly harvested embryos in PBS with heparin (10 Units/mL) (Sigma #H3393) or incubated in 1X RBC lysis solution (Roche #11814389001) for 10 minutes at room temperature, fixed at room temperature in 4% PFA for one hour, and incubated with gentle rocking in clearing solution at 37°C for 2-4 days until clear. Specimens were permeabilized and blocked for two hours to overnight with gentle rocking at 37°C in blocking buffer (PBS containing 5% normal donkey serum and 0.8% to 1.5% Triton X-100, dependent on embryo/heart stage). Specimens were labeled with primary antibody (list below) in blocking buffer with gentle rocking at 37°C, either overnight for embryos with stages up to E10.5, or for 5 days for E13.5 or E14.5 hearts. For long incubations, antibody solution was replaced halfway. Following three washes in blocking buffer, each 45 minutes except the third overnight for E13.5 and E14.5 hearts, specimens were incubated with secondary antibody (raised in donkey; AF488-, Dy405-, Cy3, or AF647-conjugated; used at 1:750; Jackson ImmunoResearch) and DAPI in blocking buffer with gentle rocking at 37°C, 2-3 hours for embryos up to E10.5, or 5 days for E13.5 or E14.5 hearts. Following three washes in PBS with gentle rocking at 37°C, samples were stored (up to overnight) in PBS with 0.2% sodium azide. Primary antibodies used were: tdTomato (rabbit, Rockland 600-401-379, 1:1000), MEF2c (sheep, R&D AF6786, 1:250), TNNT2 (mouse, Thermo MS-295-P, 1:500). Strong ZsGreen fluorescence persisted after clearing and did not require immunolabeling.

### Lightsheet Image Acquisition

Images were acquired by lightsheet microscopy, as described previously ^71^. Brielfy, specimens were warmed and transferred to 2% low melting point agarose (Fisher BP165-25) in PBS at 37°C, then embedded in glass capillary tubes with paired pistons (Sigma Z328510 paired with BR701938, or Sigma Z328502 paired with BR701934) for embryos, or tip-truncated 1mL syringes (Becton Dickinson) for hearts. After the gel solidified, the capillaries or syringes were suspended specimen-down from 14mL polystyrene tubes sealed with Parafilm. Columns containing the specimen were partially extended into an ample volume optical clearing solution (OCS, EasyIndex EI-Z1001, LifeCanvas) overnight. Following overnight incubation in OCS, specimen capillaries were retracted, and were brought to a LightSheet Z.1 microscope coupled with standard-issue laser lines and filter configurations, as well as dual pco.edge 4.2 cameras, with ZEN software (Carl Zeiss Imaging). Using immersion in OCS, specimens were imaged with a multi-view whole-volume approach, using one of two objective setups: EC Plan Neofluar 5X/0.16 with 5X/0.1 pair (hearts), or Clr Plan Neofluar 20X/1.0 paired with 10X/0.2 clearing pair (embryos). Z-stacks were collected at each view angle at the optimal slice thickness determined by Zeiss’ Zen software, ranging from 1.42 to 4.95 µm.

### Image Dataset Preprocessing

Imaging data was processed, as previously described ^71^. All data processing was performed on an 8-core x86-64 desktop PC with 64GB RAM, running Kubuntu 20.04 LTS, principally using Fiji software ^72^. Acquired lightsheet image stacks were subjected to single-view deconvolution using regularized generalized Tikhonov filtering (Parallel Spectral Deconvolution, https://imagej.net/plugins/parallel-spectral-deconvolution) following theoretical PSF calculation using the intersection of Gaussian z-plane illumination with Gibson-Lanni widefield epifluorescence detection patterns ^73^. Deconvolved stacks were co-registered, and image volumes were generated for each desired orientation, for each specimen, using Content Based Fusion in BigStitcher ^74^. TrackMate ^75^ and ImageJ 3-D viewer were used for segmentation and visualization of fused volumes.

### Image Volume Quantification

Extensive whole-volume quantifications of imaged E14.5 hearts were performed using Fiji and associated plugins, with analysis conducted with R. For assessing the location of fluorescence signal, volumes were downsampled, and 2-dimensional regions of interest (ROIs) corresponding to anatomic structures were manually drawn on each Z-slice, using the DAPI channel in a single blinded fashion. Integrated intensity for each fluorescence channel was programatically measured for each ROI, and counts were normalized to DAPI signal and pooled by anatomic region. For whole-heart quantifications, counts were further normalized to combined Ai6 plus Ai66 signal, then averaged across each anatomic region. For orientation assessments, a multi-channel panel of 174 full resolution four-chamber image excerpts, evenly distributed between apical-posterior, apical-anterior, posterior-basal, and posterior-apical IVS, were split by channel (tdTomato or ZsGreen) and blindly sampled. Using automated batch processing, these samples were subjected to automated background filtering, including contrast enhancement, and measurements of orientation coherency (OrientationJ plugin, https://github.com/Biomedical-Imaging-Group/OrientationJ), and of dominant direction and “directionality” score (Directionality plugin, https://github.com/fiji/Directionality). Directionality score distributions were rotated so that dominant direction was set to 0°, with average score (across the panel of images) was plotted as a function of direction, with testing of the distribution by the Watson U2 test (R circular package) and analyzed by Watson’s two-sample test. Individual scores surrounding 0° (±5°), 90° (±5°), and 65° (±20°) were pooled and compared with the Wilcoxon signed-rank test. An alternative measure, orientation coherency scores, across the panel of images were plotted by genotype and compared with the Wilcoxon signed-rank test.

### Quantification of Right-Left Position of Lineages

For each E10.5 heart analyzed, five representative coronal optical sections were selected, approximately 20-35 µm apart. For each optical section, the DAPI channel was used to draw five linear profiles spanning the ventricular chambers from right to left. Linear profiles were drawn such that the “0” position corresponded to the midline, identified by the interventricular groove. Fluorescence intensity of each channel (green=*Tbx5^+^* lineage, magenta=*Mef2cAHF+* lineage) was measured along each linear profile, and intensity values were normalized to the maximum value along each profile. Profiles for each channel were grouped by experimental condition (WT or DTA), and then the average ± SEM normalized fluorescence intensity profiles were calculated and plotted. Welch’s two-sample t-test was used at each position along the right-left axis to determine regions of significant difference in fluorescence intensity of each channel between experimental groups, indicating a change in the distribution of cells of that lineage.

The same images and Z-slices were also analyzed for the percent distribution of the green=*Tbx5^+^* and pink=*Mef2c-AHF^+^* lineages across the ventricles. ROIs were assigned using the DAPI channel to subset the left and the right ventricles using the IV groove as a landmark. The percent area was calculated using the area of the green or red signal and the total ventricular area. These values were normalized per Z-slice to the total area of signal in both ventricles. Significance was measured using T-tests.

For each E13.5-E14.5 heart analyzed, maximum z-projections were generated from optical sections spanning every 50 µm (E13.5-E14.0) or 100 µm (E14.5) from the anterior to posterior of the heart. For each maximum z-projection, the DAPI and red or magenta (*Tbx5*^+^/*Mef2cAHF*^+^ intersectional lineage) channels were split and background subtraction was performed on the red or magenta channel using a 50-pixel rolling ball radius. The DAPI channel was used to draw three linear profiles (basal, middle, apical) spanning the ventricular chambers from right to left. Linear profiles were drawn such that the “0” position corresponded to the middle of the IVS. Fluorescence intensity of the red or magenta channel was measured along each linear profile, and intensity values were normalized to the maximum value along each profile. Profiles were grouped by experimental condition (*Tbx5^CreERT2/+^* or *Tbx5^CreERT2/flox^*) or (e.g. WT, *Ntn1*^+/-^, *Ntn1^-/-^*) and then the average ± SEM normalized fluorescence intensity profiles were calculated and plotted. Two sample F-test and Welch’s t-test were used to analyze the distribution of *Tbx5^+^/Mef2cAHF^+^* intersectional lineage cells.

### Cell Harvesting for Single Cell RNA Sequencing

Samples for single cell RNA sequencing were collected from three independent litters at E13.5. The heart was microdissected to obtain the interventricular septum (IVS), left ventricular (LV) and right ventricular (RV) regions. Control samples were *Tbx5^CreERT2/+^;Mef2cAHF-DreERT2;ROSA26^Ai66/Ai6^* (n=3) and *Tbx5^CreERT2/+^;Mef2cAHF-DreERT2;ROSA26^Ai66^*^/+^ (n=1) for LV, RV and IVS each. *Tbx5* mutant samples were *Tbx5^CreERT2/flox^;Mef2cAHF-DreERT2;ROSA26^Ai66/Ai6^* for RV and LV (n=3 each) and IVS (n=2). One *Tbx5* mutant sample of the IVS+AV complex regions was not included because it was lost from a microfluidic chip clog during sample processing.

Each microdissected tissue was singularized with TrypLE Express (Life Technologies, Cat# 12604-013) at 37C and quenched with 1% FBS in PBS. The single cell suspension was then filtered through a cell strainer cap (Corning, Cat# 352235) and centrifuged at 300g for 5 minutes. The pellet was resuspended in 1% FBS in PBS and counted using an automated cell counter. A 30μL aliquot of the cell suspension was used to generate single cell droplet libraries with the Chromium Next GEM Single Cell 5’ Library & Gel Bead Kit v1.1, according to manufacturer’s instructions (10X Genomics). After KAPA qPCR quantification, a shallow sequencing run was performed on a NextSeq 500 (Illumina) prior to deep sequencing on a NovaSeq S4 (Illumina). For control IVS+AV complex regions, at E13.5, 3 samples were *Tbx5^CreERT2/+;^Mef2cAHF-DreERT2;ROSA26^Ai66/Ai6^*

### Data Processing Using Cellranger

All datasets were processed using Cellranger 2.0.2. FASTQ files were generated using the mkfastq function. Reads were aligned to a custom mm9 reference (version 1.2.0) containing tdTomato and ZsGreen reporter genes. Cellranger aggr was used to aggregate individual libraries after read depth normalization.

### Seurat Analysis

Outputs from the Cellranger pipeline were analyzed using the Seurat v3 ^76–78^. Datasets from the IVS, LV and RV were analyzed separately. Cells with 10,000-50,000 UMIs and 1500-7250 genes were retained. Data was normalized using the NormalizeData function and scaled using ScaleData, while also regressing unwanted sources of variation, such as differences in the number of UMIs, number of genes, percentage of mitochondrial reads and differences between G2M and S phase scores. Principal component analysis (PCA) was performed using the most highly variable genes. Cells were then clustered based on the top 30 principal components and visualized on a Uniform Manifold Approximation and Projection (UMAP) ^79^. A clustering resolution that separated cells by major cell types was chosen. Next, *Tnnt2^+^* clusters from the IVS dataset were extracted and re-clustered, similar to the parent dataset above. A clustering resolution was chosen based on separation of genotypes. Differential gene expression tests between clusters or between *ZsGreen*^+^ and *ZsGreen*^-^ or *tdTomato*^+^ and *tdTomato*^-^ cells were performed using the FindMarkers function with default parameters. Selected differentially expressed genes with an adjusted p-value less than 0.05 from the Wilcoxon rank sum test were displayed using the Dotplot or FeaturePlot functions.

### Fluorescent *In Situ* Hybridization

E11.5 or E14.5 hearts were fixed with 4% paraformaldehyde or 10% formalin overnight at 4C, embedded in paraffin and then sectioned for a transverse or four-chambered view on slides. *In situ* hybridization was performed on sections using the RNAscope Multiplex Fluorescent v2 Assay kit (Advanced Cell Diagnostics, Cat# 323100). Briefly, sections were deparaffinized in xylene and 100% ethanol, treated with hydrogen peroxide for 10 minutes and boiled in target retrieval buffer for 10 minutes. A hydrophobic barrier was drawn around each section using an Immedge pen (Vector Laboratories, Cat# H-4000), and slides were allowed to dry overnight. The following day, sections were treated with Protease Plus for 30 minutes, followed by hybridization with probes for 2 hours at 40C. Probes (Advanced Cell Diagnostics) used were *Mm-Tnnt2-C4* (Cat# 418681-C4), *Mm-Ntn1* (Cat# 407621), *Mm-Slit2* (Cat# 449691), *Mm-Nppa-C3* (Cat# 418691-C3), *Mm-Bmp2-C2* (Cat# 406661-C2), *Mm-Robo1* (Cat#475951), *Mm-Unc5b* (Cat#482481), and *Mm-Irx2* (Cat#519901). Amplification steps were carried out according to manufacturer’s instructions. Opal dyes 520, 570, 620 and 690 (Akoya Biosciences, Cat# FP1487001KT, FP1487001KT, FP1487001KT and FP1487001KT) were used at 1:750 dilution. After staining with DAPI, 1-2 drops of ProLong Gold Antifade Mountant (ThermoFisher Scientific, Cat# P36930) were placed on the slides and mounted with a coverslip. Slides were stored overnight at 4C and were imaged at 10X magnification on the Olympus FV3000RS. Multi-Area Time Lapse (MATL) images were captured and then stitched together using the Olympus FV3000RS software. Stitched images were analyzed using ImageJ OlympusViewer plugin.

### Immunohistochemistry

Cryopreserved slides were thawed from −80C at RT and washed in 0.1% Triton X-100 in PBS (PBST) for 3 x 5 min. Antigen-retrieval was performed by boiling slides in 10mM sodium citrate buffer (pH 6.0) for 10 minutes and then washed in H2O for 3 x 5 min. Blocking was performed using 5% donkey serum in 0.1% PBST at RT for 1-2hr. Anti-Netrin-1 antibody 1:500 (R&D Systems AF1109) in 1% BSA + 0.1% PBST was incubated at 4C overnight. After washing 3 x 5 min in 0.1% PBST, slides incubated in Alexa Fluor 594 (1:300) in 1% BSA + 0.1% PBST at RT for 1 hr. Slides were then washed 3 x 5 min in 0.1% PBST and incubated in 1:1000 DAPI in 0.1% PBST at RT for 5-10 minutes. Finally, slides washed 3 x 5 min in 0.1% PBST, glass coverslips were mounted using Prolong Gold antifade mounting medium and stored at 4C.

### Quantification of IVS Cell Size and Shape by WGA Staining

Blind measurements of interventricular septum (IVS) cell size and shape were made on 5 μm paraffin-embedded *Tbx5*^CreERT2/+^ or *Tbx5*^CreERT2/flox^ hearts at E14.5, sectioned in 4-chamber view approximately midway in the anterior-posterior axis. After sectioning, slides were stained for cardiac troponin T (cTNT, 1:500 cTNT, Thermo, MS-295-P) using Alexa 488 donkey anti-mouse secondary antibody (Jackson Immunoresearch), in addition to Alexa 594 conjugated Wheat Germ Agglutinin (WGA, dilution 5 μM, Thermo, W11262) and DAPI. After widefield imaging on a confocal fluorescence microscope at 40X, regions of interest (ROIs) from apical, basal, and mid regions of the IVS were extracted in both DAPI and WGA channels, guided by the cTNT staining as an anatomic reference. ROIs from the DAPI channel were subjected to automatic segmentation and counting to assess cell number and density. ROIs from the WGA channel were scrambled for blinding *Tbx5* genotype, and 24 to 56 easily identified cell bodies per ROI (depending on ROI size) were manually assessed in Fiji for length, width, and axis orientation. Taking cell bodies as roughly elliptical, their diameter, area, and eccentricities were estimated, and *Tbx5* genotype identities were unscrambled for plotting. Scripts for the filename scrambling/unscrambling, as well as automated quantification and count tallying, as described above, are available at Github (https://github.com/mhdominguez/Kathiriya-NCVR-2025-analysis).

### Machine Learning-Based 3-D Reconstruction from Histology and Quantitative Micrometry

Tissues were formalin-fixed, paraffin embedded (FFPE) and serially sectioned at a standard thickness of 5 µm to exhaustively collect the heart tissue of each fetal mouse. Serial sections were stained with hematoxylin and eosin (H&E) and digitized at 20x magnification using a Leica Versa 200 Automated Slide Scanner. CODA, a technique to 3-D reconstruct serial tissue images, was used to map the microanatomy of the hearts ^58,80^. The various steps of CODA can be broken down into image registration, nuclear detection, and tissue labelling. Nonlinear image registration was used to align the serial images. Cellular coordinates were generated through color deconvolution and detection of two-dimensional intensity peaks in the hematoxylin channel of the images. A deep learning semantic segmentation algorithm was trained to label the anatomical structures of the mice at 1 µm resolution. To train the model, manual annotations of six structures were generated in 55 histological images corresponding to the different large structures present in the tissue: bones, spinal cord, lung, liver, stroma, heart, and non-tissue pixels in the images. A second deep learning model was trained to subclassify the regions of interest within the heart: compact myocardium, trabeculae, IVS, AV complex region, and atria. Fifty images from the manual annotation datasets were used for model training, with five images held out for independent testing of model accuracy. The models were deemed acceptable when they reached a minimum per-class precision and recall of >85%. Using the trained model, independent images were segmented and aligned to generate digital 3-D datasets.

The reconstructed volumes enabled microanatomical quantification of anatomical properties. IVS fill, or the solidity of the muscle layer in the IVS, was calculated by dividing the number of dark (<200 in RBG color space) pixels within the IVS, normalized by all pixels (including empty space) within the IVS. Percent trabeculation within the IVS was calculated by isolating the tissue classified as trabeculae that was spatially attached to the IVS, then normalizing this number by the combined volume of trabeculae attached to IVS and the volume of the IVS itself. The compact myocardium thickness was calculated by measuring the thickness of the compact layer at the bottom 20% of the ventricles. To measure the thickness on the left and right ventricles, the two sides of the heart were manually segmented in 3-D space. VSDs were identified by calculating regions where the empty space of the left ventricle contacted the empty space of the right ventricle. These defects were evaluated and manually sorted into membranous or muscular sub-types. The areas of each defect were then manually calculated through annotation on serial histological images representing the shortest path to close the defect. These lines were summed across all images containing the defect multiplied by the thickness of the histological slides to determine the area of the hole.

### Statistics and Reproducibility

For statistical analysis of genetic interactions, the number of animals with a given defect (muscular VSD or membranous VSD) was modeled in a Generalized Linear Model assuming a binomial probability distribution for the observed counts. This model included main effects (in terms of log odds ratios) capturing the number of mutant alleles for *Tbx5* (0 or 1, representing the WT or the Het genotype, respectively) and the number of mutant alleles for *Slit2* or *Ntn1* (0,1 or 2: representing the WT, Het or Hom genotype, respectively) and an interaction between these effects. Given the relatively small number of animals in this experiment, bias-reduced estimates were made using the brglm2 package {Kosmidis.2020} in R.

Representative images are shown based on experiments that were repeated independently with similar results, as follows: Figure(s) 1a-f: 2 samples; 1g-j: 5 samples; 2a-i: 4 samples; 2j-n: 6 samples; 2o-q: 8 samples; 2r-t: 15 samples; 3a,b: 2 controls, 2 *DTA* mutants; 3c-j: 2 controls, 4 *DTA* mutants; 3l-n’ and Extended Data 2: 3 controls, 2 *DTA* mutants; 4a-d: 5 controls and 4 *Tbx5* mutants, for which some samples are shown in Extended Data Figure 4a-i; Figure 4i-m: 4 controls, 4n-r: 3 *Tbx5* mutants; 4u-v’; 2 controls, 2 *Tbx5* mutants; 5i-n; 2 controls and 2 *Tbx5* mutants, for which the second replicates are shown in Extended Data Figure 6i-n”; Figure 6f-h: 14 controls, 8 *Slit2*^+/-^, 4 *Slit2*^-/-^; 6j-l: 14 controls, 9 *Ntn1^+/-^,* 7 *Ntn1*^-/-^, with some samples shown in Extended Data Figure 10a-d; Extended Data Figure(s) 1: 2 samples per condition; 4a-l: 5 controls, 4 *Tbx5* mutants, with some samples shown in Figure 4; 3: 3 controls, 7 *Tbx5* mutants; 6a-h’: 2 controls and 2 *Tbx5* mutants; 6o: 2 controls; 7a-h: 2 controls and 2 *Tbx5* mutants; 9a-d: 14 controls, 8 *Tbx5*^+/-^, 8 *Slit2*^+/-^; 8 *Tbx5*^+/-^;*Slit2*^+/-^; 9f-i: 14 controls, 7 *Tbx5*^+/-^; 9 *Ntn1*^+/-^, 7 *Ntn1*^-/-^. Supplemental Figure(s) 1: 5 samples; 2: 5 controls, 4 *Tbx5* mutants; 3: 2 controls, 2 *Tbx5* mutants.

## Data Availability

Source data is available upon publication. scRNA-seq data generated in this paper is available in the NCBI GEO database (GSE260601). ChIP-seq data from Akerberg et al., 2019, is available in the GEO database (GSE124008).

## Code Availability

Code related to “Image Volume Quantification” ^71^ is available at https://github.com/mhdominguez/LSFMProcessing. Source data and scripts used to analyze other morphometrics from lightsheet microscopy, such as right-left positioning, orientation and directionality of cells, are available at https://github.com/mhdominguez/Kathiriya-NCVR-2025-analysis.

## Acknowledgements

We dedicate this manuscript in memory of Tatyana Sukonnik (Gladstone Institutes). We thank Brian Black (University of California, San Francisco) for the *Mef2cAHF-Cre* mouse line, and members of the Bruneau lab for troubleshooting help, discussions and comments. We also thank the Gladstone Bioinformatics, Genomics, and Histology and Microscopy Cores, the UCSF Laboratory for Cell Analysis, and the UCSF Center for Advanced Technology. This work was supported by grants from the National Institutes of Health (NHLBI Bench to Bassinet Program UM1HL098179 to B.G.B.; R01HL114948 to B.G.B., R01HL155906 to B.G.B. and I.S.K., 1R56HL166894-01A1 to I.S.K., K99HL177318 to J.M.M-V, U54CA268083 and UG3CA275681 to A.L.K.), also by a postdoctoral fellowship from the American Heart Association, together with The Children’s Heart Foundation (24POST1191660) to J.M.M.-V., Additional Ventures Innovation Fund to B.G.B., Society for Pediatric Anesthesia (Young Investigator Award to I.S.K.), Hellman Family Fund (I.S.K.), UCSF REAC Grant (I.S.K.), UCSF Pediatric Heart Center Catalyst Award to I.S.K. and UCSF Department of Anesthesia and Perioperative Care Research Support to I.S.K. This work was also supported by an NIH/NCRR grant (C06 RR018928) to the J. David Gladstone Institutes, and the Younger Family Fund (B.G.B.).

## Author Contributions Statement

W.P.D. and B.G.B. originally conceived, and I.S.K. and B.G.B. refined and developed the study. W.P.D. made initial observations using genetic lineage tools that he gathered or generated. I.S.K., M.H.D., W.P.D. and K.M.H. performed embryo dissections. K.M.H., B.I.G., P.G., M.N.M., T.S., D.M.P., S.W. and E.F.B. managed animal husbandry and genotyping. I.S.K., M.H.D., W.P.D. and K.M.H. performed whole mount or section immunostaining experiments and imaging. M.H.D., J.M.M. and D.Q performed lightsheet imaging and processing, and M.H.D. and J.M.M. implemented analyses with hypothesis-based input from I.S.K. I.S.K. designed the strategy to use 5’ sequencing kit and developed synthetic genome with Gladstone Bioinformatics core, micro-dissected heart regions for scRNA-seq and processed samples with K.S.R. and P.G. for sequencing library preparations. K.S.R. generated scRNA-seq libraries and performed scRNA-seq analyses with hypothesis-based input from I.S.K. K.S.R., K.M.H., P.G. and M.N.M. performed section *in situ* hybridization and imaging. K.S.R. generated ChIP-seq browser tracks with input from S.K.H. I.S.K. assessed cardiac defects by histology and provided anatomical knowledge to A.L.K. to apply a deep learning tool for cardiac morphometry. A.F. trained deep learning models and was supervised by A.L.K. and advised by P-H.W. and D.W. B.G.B. supervised and advised the study. I.S.K. and K.S.R. prepared figures. I.S.K. wrote the original draft with contributions from co-authors. I.S.K. and B.G.B. edited the manuscript with input from co-authors.

---

**Supplementary Figure 1.**
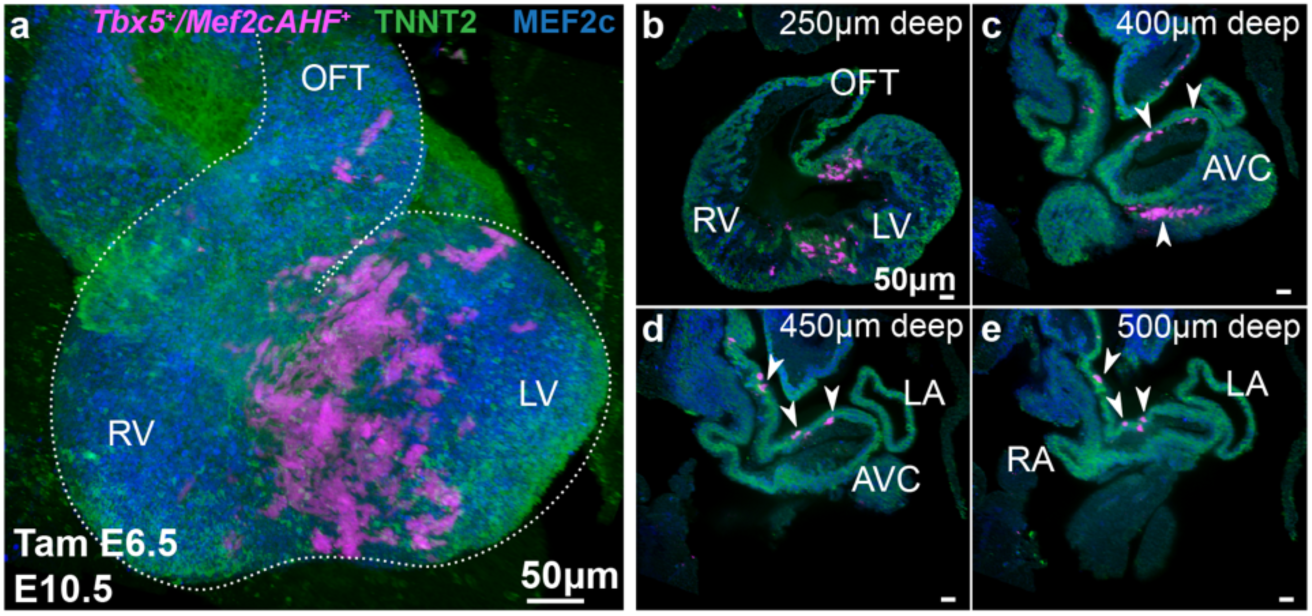
*Tbx5^+^/Mef2cAHF^+^* lineage contributions to the developing atria. **a-e**, In the developing atria at E10.5, optical sections from ventral to dorsal show lineage-labeled *Tbx5^+^/Mef2cAHF^+^* lineage cells marked at E6.5 by a single tamoxifen dose is located from the AV canal (AVC) region to the roof of the atrium, prior to the formation of the interatrial septum, in *Tbx5^CreERT2^*^/+^;*Mef2cAHF-DreERT2*;*ROSA26^Ai6^*^/*Ai66*^ embryos. All scale bars equal 50 microns. **Video 1.** Related to Figure 2 and Supplemental Figure 1.

**Supplemental Figure 2.**
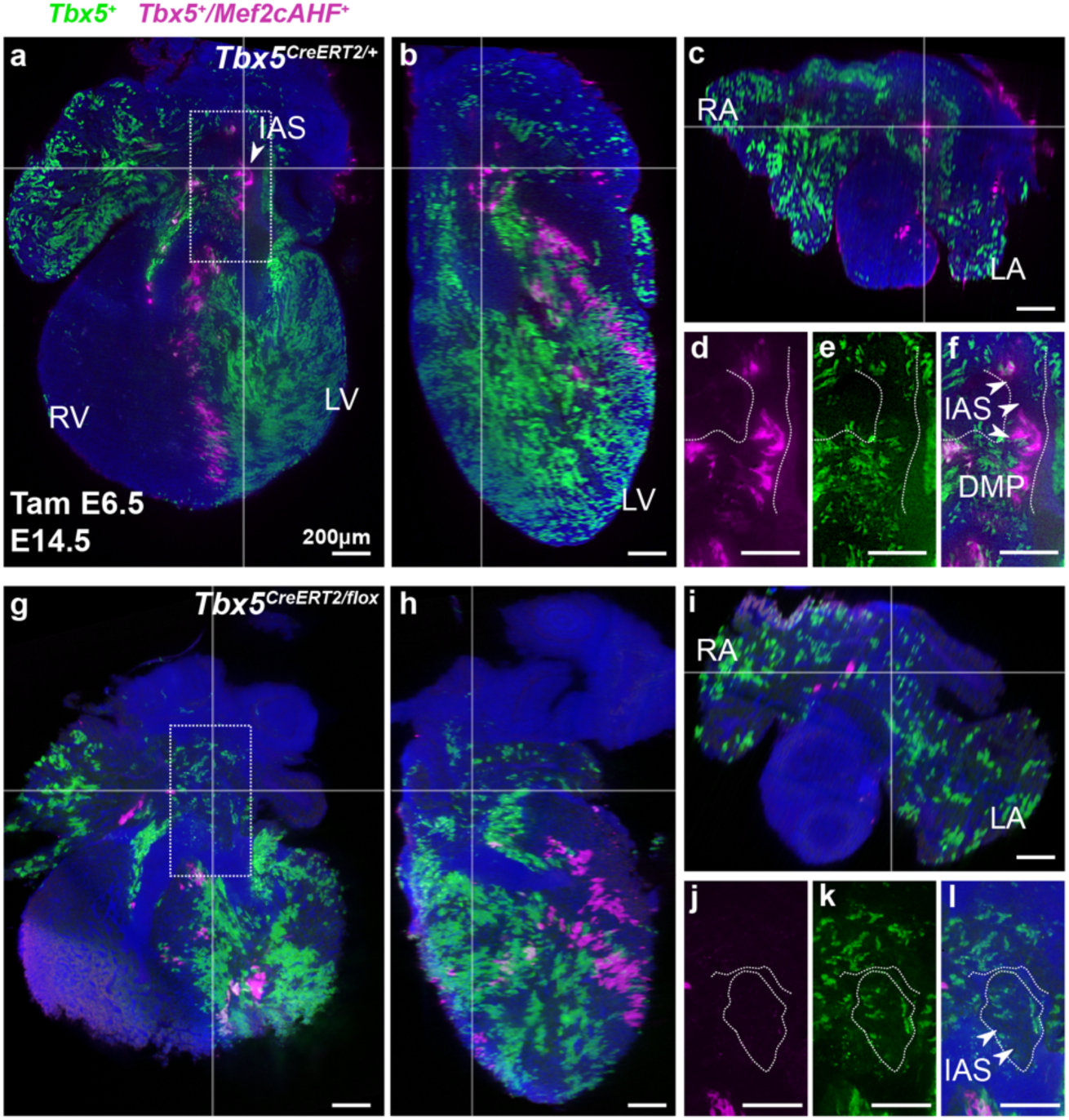
Reduced TBX5 dosage leads to reduced contributions of the *Tbx5^+^/Mef2cAHF^+^* lineage to the inter-atrial septum and dorsal mesenchymal protrusion. **a-f,** At E14.5, the *Tbx5*^+^/*Mef2cAHF*^+^ lineage cells contribute to the interatrial septum (IAS, arrowheads) and dorsal mesenchymal protrusion (DMP) in *Tbx5^CreERT2/+^* controls (*Tbx5^CreERT2/+^*;*Mef2cAHF-DreERT2*;*ROSA26^Ai6/Ai66^*). **g-l**, The *Tbx5*^+^/*Mef2cAHF*^+^ lineage cells are less apparent in *Tbx5* mutants (*Tbx5^CreERT2^*^/*flox*^;*Mef2cAHF-DreERT2*;*ROSA26^Ai6^*^/*Ai66*^). Right ventricle (RV), left ventricle (LV), right atrium (RA), left atrium (LA). All scale bars equal 200 microns.

**Supplemental Figure 3.**
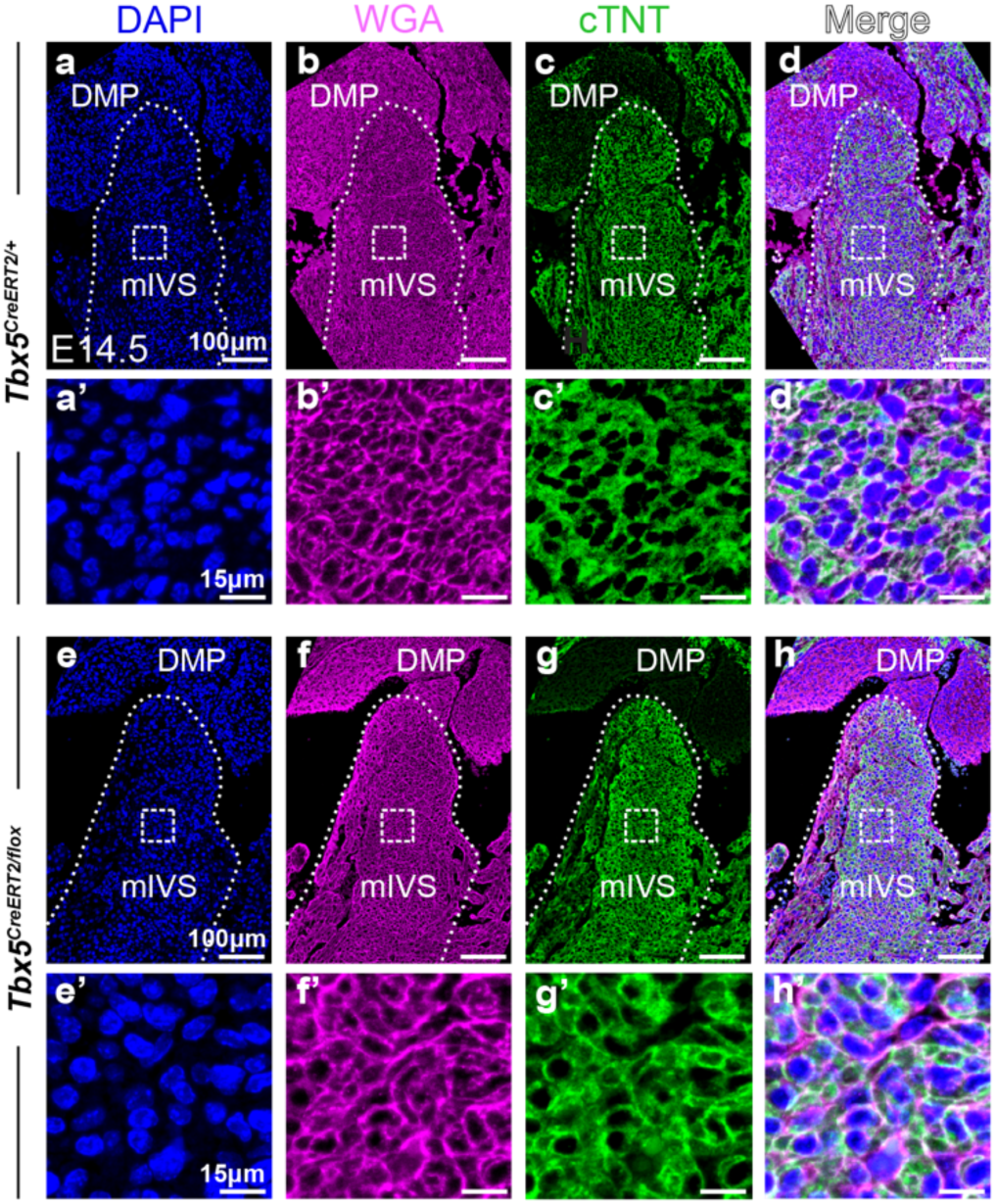
Images for quantification of cell geometry, and assessment of tissue architecture in *Tbx5* mutants. **a-d’**, *Tbx5^CreERT2/+^* control (*Tbx5^CreERT2/+^*;*Mef2cAHF-DreERT2*;*ROSA26^Ai6/Ai66^*) (3 planes per sample, 2 samples) and (**e-h’**) *Tbx5^CreERT2/flox^* mutant (*Tbx5^CreERT2/flox^;Mef2cAHF-DreERT2*;*ROSA26^Ai6/^*^Ai66^) hearts (3 planes per sample, 2 samples) after single dose of tamoxifen injected at E6.5 is shown at E14.5, stained with DAPI, wheat germ agglutinin (WGA), or cardiac troponin T (cTNT). (b, b’, f, f’) are repeat images of Figure 4u-v’. scale bars: 100 microns (a-h); 15 microns (a’-h’).

**Supplementary Figure 4.**
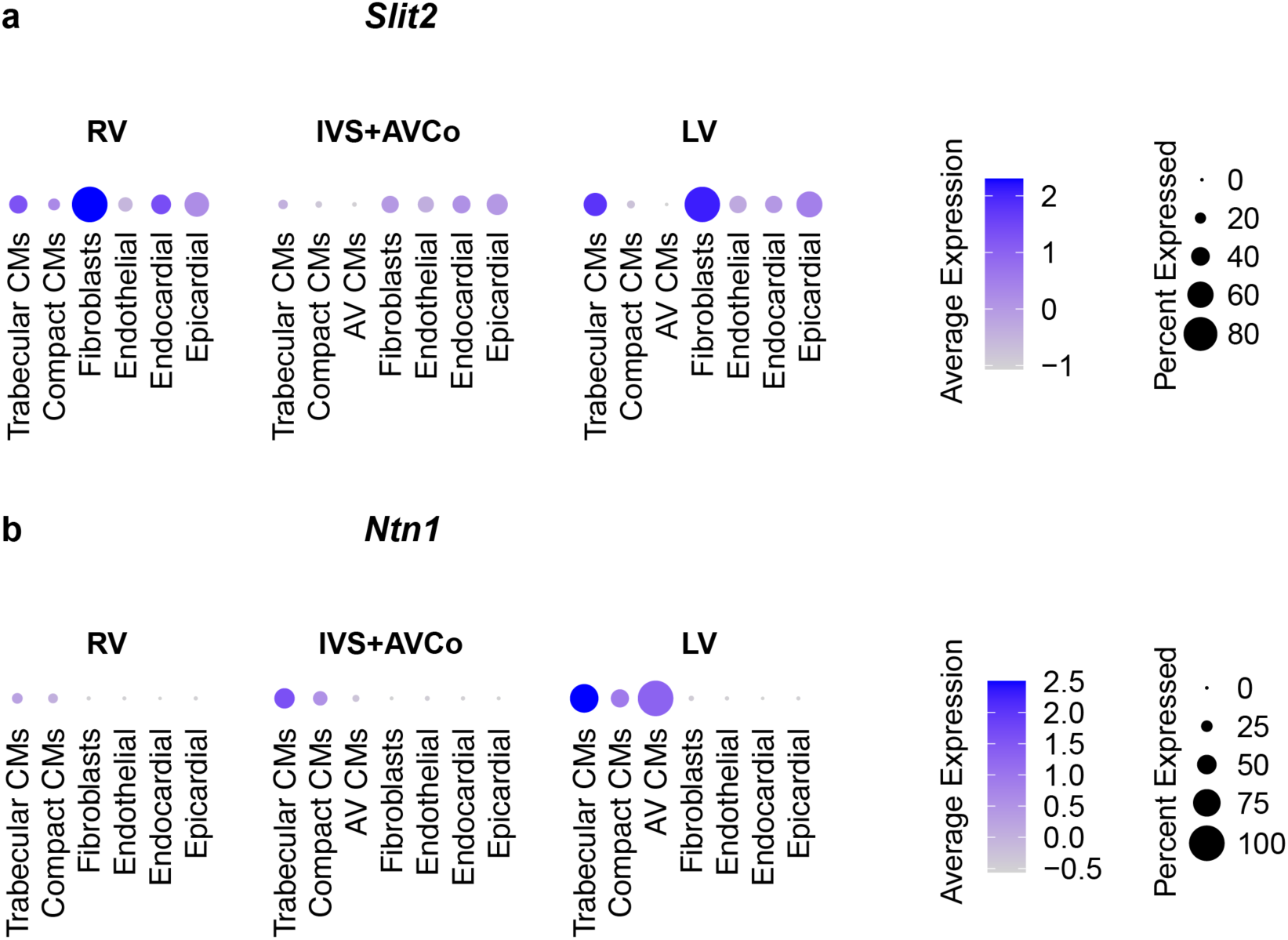
Gene expression of *Slit2* and *Ntn1* by cell type in individual cells of the developing heart. By scRNA-seq from control E13.5 hearts (n=4), gene expression of (**a**) *Slit2* and (**b**) *Ntn1* are shown by region (left ventricle (LV), right ventricle (RV), interventricular septum (IVS)+atrioventricular complex (AVCo) regions).

**Extended Data Figure 1.**
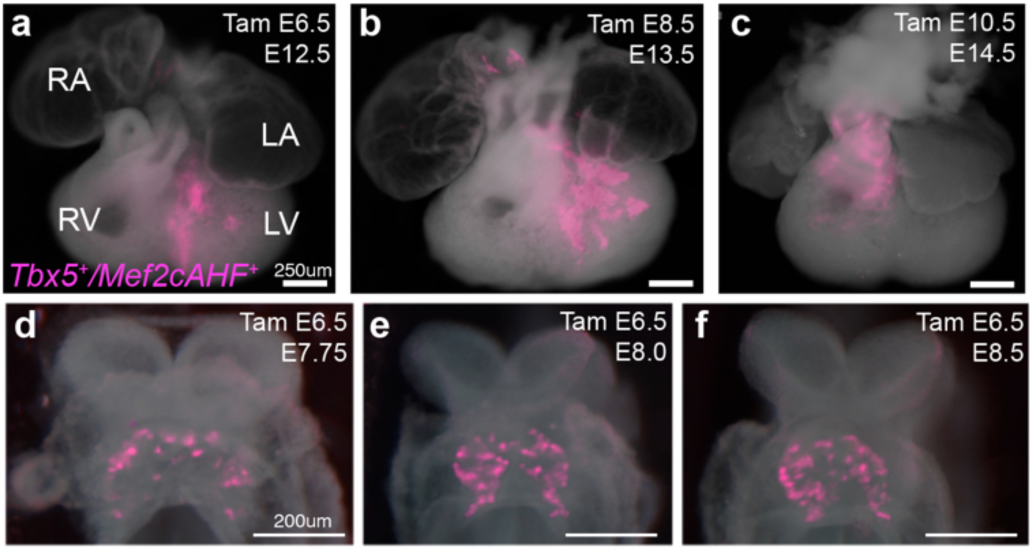
**a-c**, *Tbx5^+^/Mef2cAHF^+^* lineage contributions to the interventricular septum were descendants from progenitors labeled at (a) E6.5, but not at (b) E8.5 or (c) E10.5 in *Tbx5^CreERT2^*^/+^;*Mef2cAHF-DreERT2*;*ROSA26^Ai6^*^/*Ai66*^ embryos. Scale bars: 250 microns. **d-e**, Epifluorescence microscopy images of the *Tbx5^+^/Mef2cAHF^+^* lineage (tdTomato immunostaining) labeled by a single dose of tamoxifen at E6.5 of embryos at (d) E7.75, (e) E8.0 and (f) E8.5 displayed an apparent “salt and pepper” pattern in the early developing heart. Scale bars: 200 microns.

**Extended Data Figure 2.**
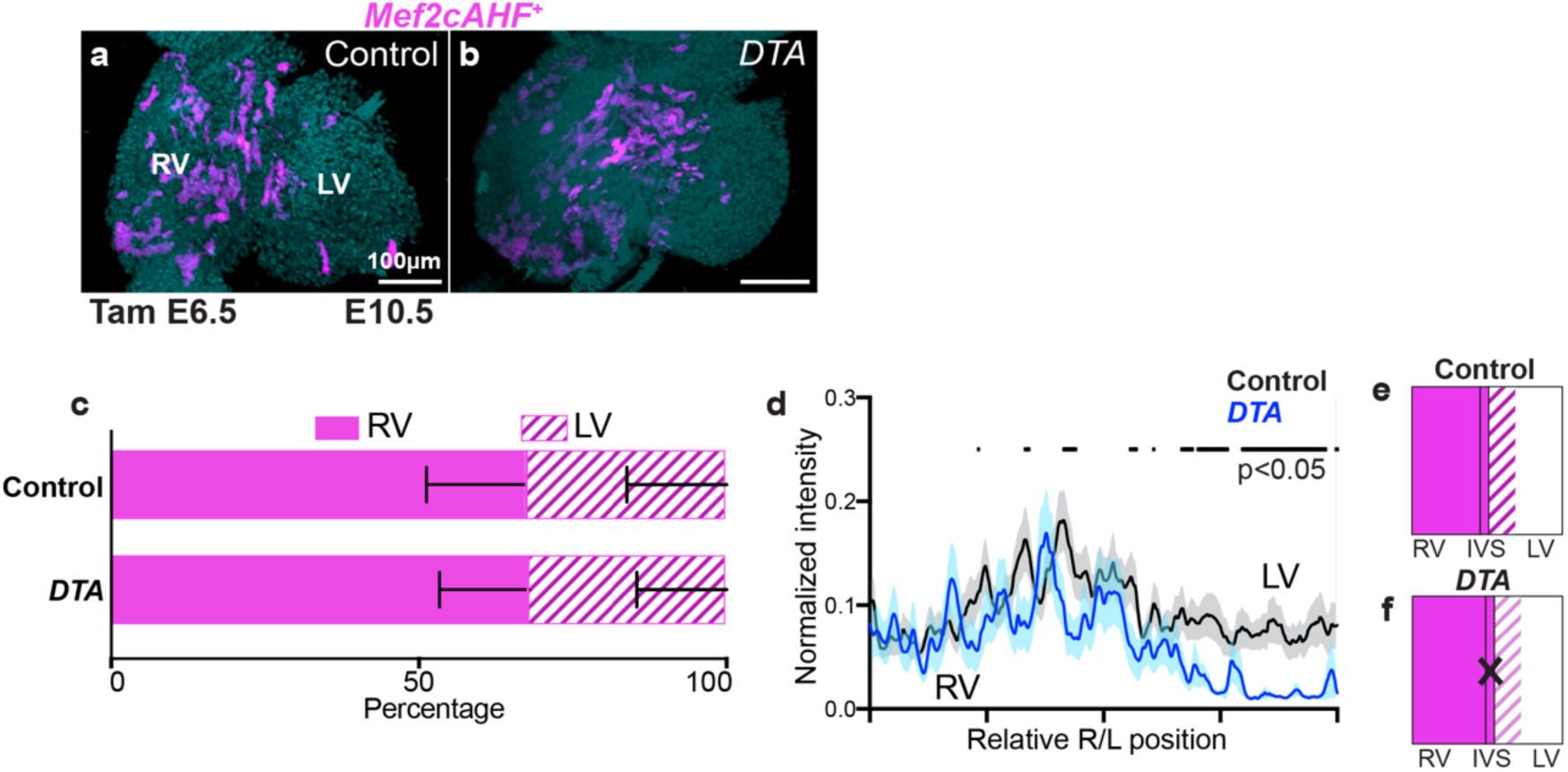
*Mef2cAHF*^+^ lineage distribution in intersectional-*DTA* mutants. **a**, *Mef2cAHF*^+^ lineage was enriched in the RV of control embryos (*Tbx5^CreERT2/+^;Mef2cAHF-DreERT2;Hip11^+/+^*) (n=3) and (**b**) intersectional-*DTA* mutant embryos (*Tbx5^CreERT2/+^;Mef2cAHF-DreERT2;Hip11^Intersectional-DTA176/+^*) (n=2). **c**, This distribution of the *Mef2cAHF*^+^ lineage was not different in the RV or LV between control and *DTA* mutants (p=0.939327, by two-sided T-test). Cells of the right ventricle (RV) or left ventricle (LV) in five optical slices at different anterior-posterior planes per embryonic heart sample of 3 controls and 2 mutants were evaluated. Mean and standard deviation are shown. **d**, Linear profiles showed a reduction of fluorescence activity in the LV. Linear profiles at 5 apical-basal planes per optical slice, for 5 optical slices at different anterior-posterior planes per embryonic heart sample of 3 controls and 2 *DTA* mutants were assessed. Statistical significance (p<0.05) between linear profiles of control and *DTA* mutants was determined by Welch’s two-sided t-test at each position along the right-left axis. Mean and standard error of the mean are shown. Precise p-values can be found in the Source data. **e, f**, Cartoon depictions.

**Extended Data Figure 3.**
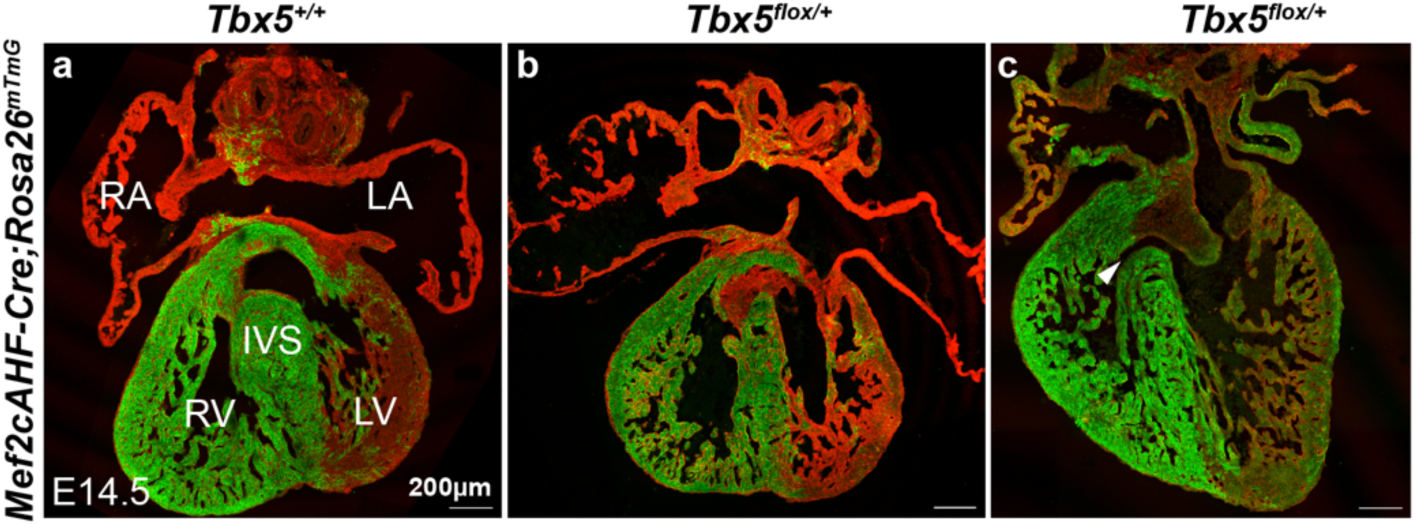
Conditional haploinsufficiency of *Tbx5* in the interventricular septum (IVS) leads to ventricular septal defects (VSDs). **a-c**, At E14.5, membranous VSDs were observed in *Mef2cAHF-Cre*;*ROSA26^mTmG/+^;Tbx5^flox/+^* mutant embryos (n=4/7) compared to *Mef2cAHF-Cre*;*ROSA26^mTmG/+^;Tbx5^flox/+^* controls (n=0/3). Right ventricle (RV), left ventricle (LV) and interventricular septum (IVS). All scale bars equal 200 microns.

**Extended Data Figure 4.**
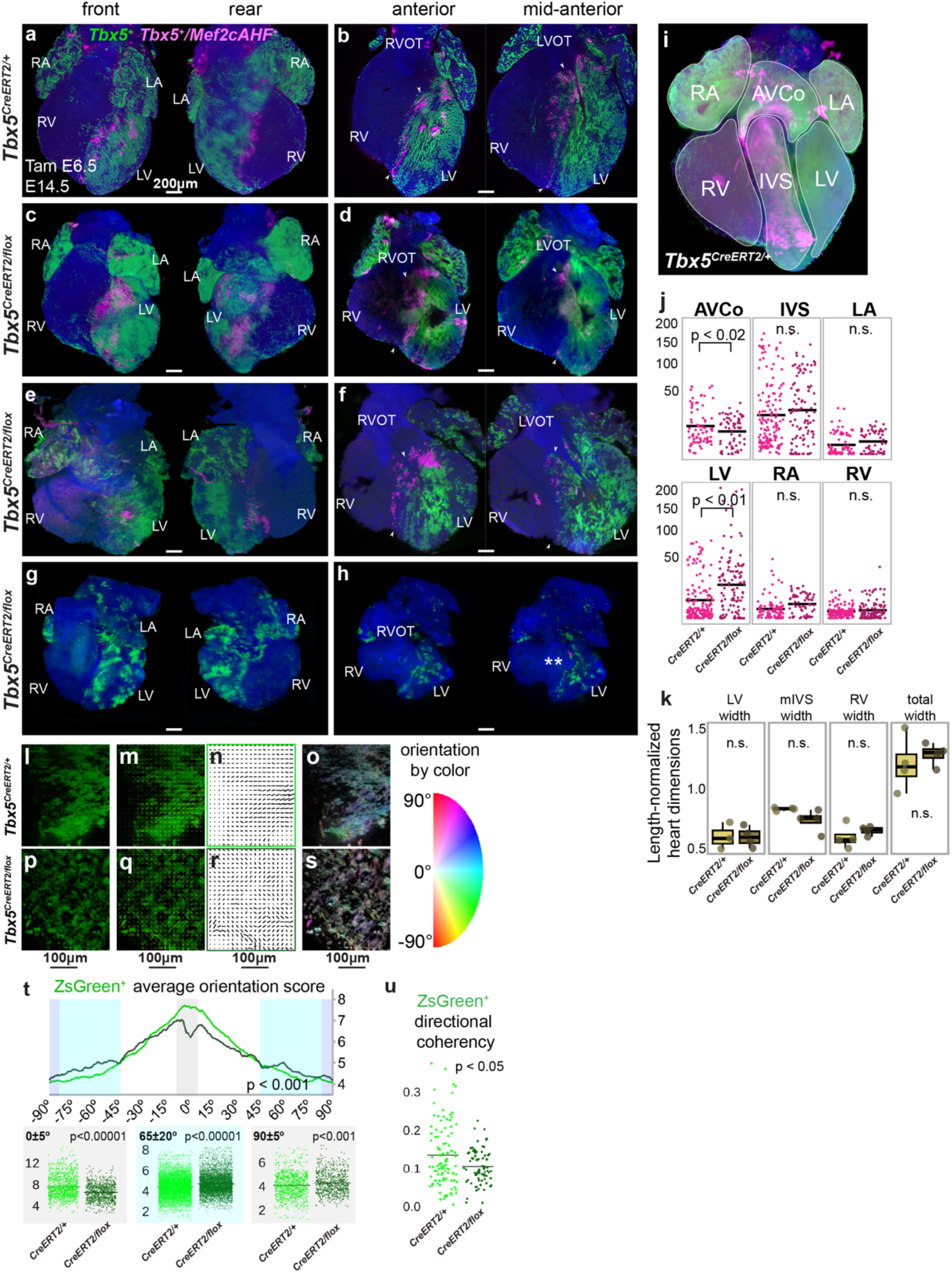
Additional samples of *Tbx5* mutant hearts and supplementary quantification of distribution and cell arrangement. **a-h**, Volume renderings from lightsheet microscopy of anterior and posterior surfaces, as well as anterior and mid-anterior 4-chamber optical sections of *Tbx5^CreERT2/+^* control (*Tbx5^CreERT2/+^*;*Mef2cAHF-DreERT2*;*ROSA26^Ai6/Ai66^*) and *Tbx5^CreERT2/flox^* mutant (*Tbx5^CreERT2/flox^;Mef2cAHF-DreERT2*;*ROSA26^Ai6/^*^Ai66^) hearts at E14.5 display *Tbx5^+^* lineage (ZsGreen) and *Tbx5^+^/Mef2cAHF^+^* lineage (tdTomato immunostaining) cells. (a-h) scale bars: 200 microns. **i-k**, Quantification of the anatomical location of tdTomato^+^ and ZsGreen^+^ cells in *Tbx5^CreERT2/flox^* mutants (n=3) and *Tbx5^CreERT2/+^* controls (n=4), by region at E14.5 as outlined in (i). Over 1200 3-dimensional regions of interest were drawn blinded, and then integrated fluorescence signal was assessed. **j**, Although no regional differences in the intensities of ZsGreen^+^ were detected, tdTomato^+^ displayed higher integrated fluorescence in the left ventricle (LV) and less signal in atrioventricular complex (AVCo) complex region by two-sided unpaired T-tests (AVCo: p=0.01933, interventricular septum (IVS): p=0.3679, left atrium (LA): p=0.2829, LV: p=0.0001574, right atrium (RA): p=0.1678, right ventricle (RV): p=0.05063). **k**, Length-normalized widths of LV, muscular IVS (mIVS), RV and total heart at E14.5 were not significantly different between control (n=4) and *Tbx5* mutants (n=3) based on regions of interest in (i), by two-sided unpaired T-tests (p=0.9654, 0.1277, 0.3797, 0.5631 for left-to-right plots). Median, first and third quartiles are shown in box plots. Further details are in Source data. **l-u**, *Tbx5^CreERT2/flox^* mutant hearts at E14.5 scored worse for orientation scores of *Tbx5^+^* lineage (ZsGreen^+^) cells in the dominant direction (range from −5 to 5 degrees) and scored higher in the orientation orthogonal to the dominant direction (range from 85 to 95 degrees and −85 to −95 degrees), as determined (**t**) by two-sided Watson U2 test across all orientations (p<0.001) or by two-sided Wilcoxon rank sum test with continuity correction for selected orientations (p-values < 2.2e-16 for left and middle plots, p-value = 0.0006863 for right plots). **u**, *Tbx5^+^* lineage (ZsGreen^+^) cells demonstrated worse directional coherency scoring in *Tbx5^CreERT2/flox^* mutants by two-sided Wilcoxon rank sum test with continuity correction (p=0.03316). (l-s) scale bars are 100 microns. **Video 2.** Related to Figure 4 and Extended Data Figure 4.

**Extended Data Figure 5.**
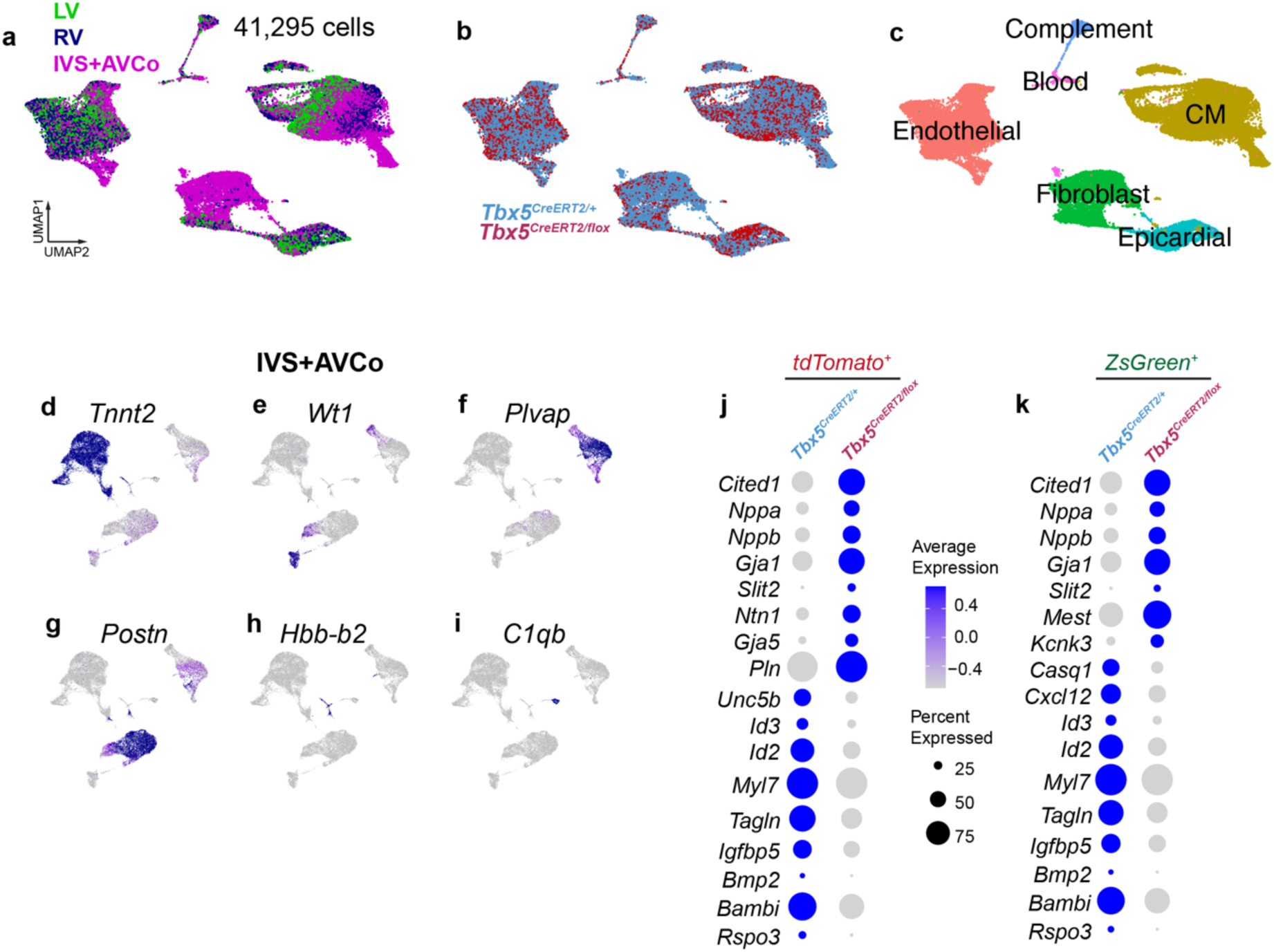
scRNA-seq of *Tbx5* mutants by region. **a-c**, In scRNA-seq samples from E13.5 hearts, we display cells by (**a**) region (left ventricle (LV), right ventricle (RV), interventricular septu (IVS)) (**b**) genotype, or (**c**) cell types by Louvain clustering. In IVS+atrioventricular complex (AVCo) regions (control, n=4, *Tbx5* mutant, n=2), we detected clusters enriched for (**d**) *Tnnt2*^+^ cardiomyocytes (CMs), (**e**) *Tbx18^+^/Wt1^+^* epicardial cells, (**f**) *Plvap*^+^ endothelial cells, (**g**) *Postn*^+^ fibroblasts, (**h**) *Hbb-b2*^+^ red blood cells, and (**i**) *C1qb^+^* white blood cells. Dot plots of selected differentially expressed genes in (**j**) *tdTomato*^+^ or (**k**) *ZsGreen*^+^ cells from the IVS+AVCo region. Significance of adj p<0.05 was determined by two-sided Wilcoxan rank sum.

**Extended Data Figure 6.**
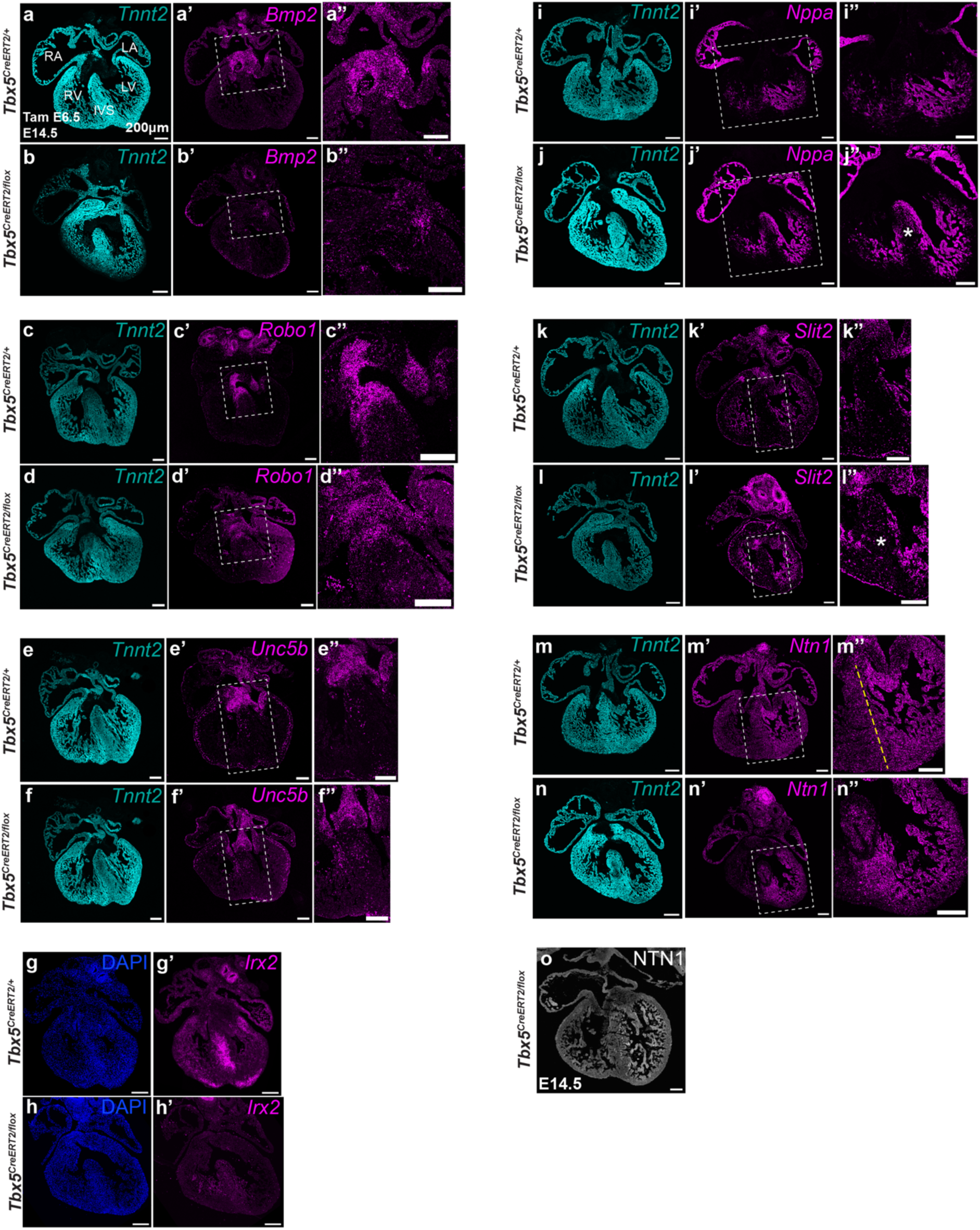
Fluorescent *in situ* hybridization of *Tbx5*-dependent genes at E14.5. **a-f”**, By fluorescent *in situ* hybridization at E14.5 after a single dose of tamoxifen at E6.5, *Bmp2*, *Robo1* and *Unc5b* are expressed in the atrioventricular canal (boxed region) in *Tbx5^CreERT2/+^* controls (*Tbx5^CreERT2/+^*;*Mef2cAHF-DreERT2*;*ROSA26^Ai6/Ai66^*), and expression of these genes are reduced in *Tbx5^CreERT2/flox^* mutants (*Tbx5^CreERT2/flox^;Mef2cAHF-DreERT2*;*ROSA26^Ai6/^*^Ai66^). *Tnnt2* is expressed in cardiomyocytes. **g-h’**, *Irx2* is reduced in the IVS of *Tbx5* mutants. **i-n”**, *Nppa*, *Slit2* and *Ntn1* are expressed in the trabecular layer in *Tbx5^CreERT2/+^* controls, and misexpressed in the IVS in *Tbx5^CreERT2/flox^* mutants with a membranous VSD at E14.5. Yellow dashed line demarcates gradient of *Ntn1* expression across the IVS from left to right. **o**, Immunostaining of NTN1 at E14.5. All scale bars are 200 microns.

**Extended Data Figure 7.**
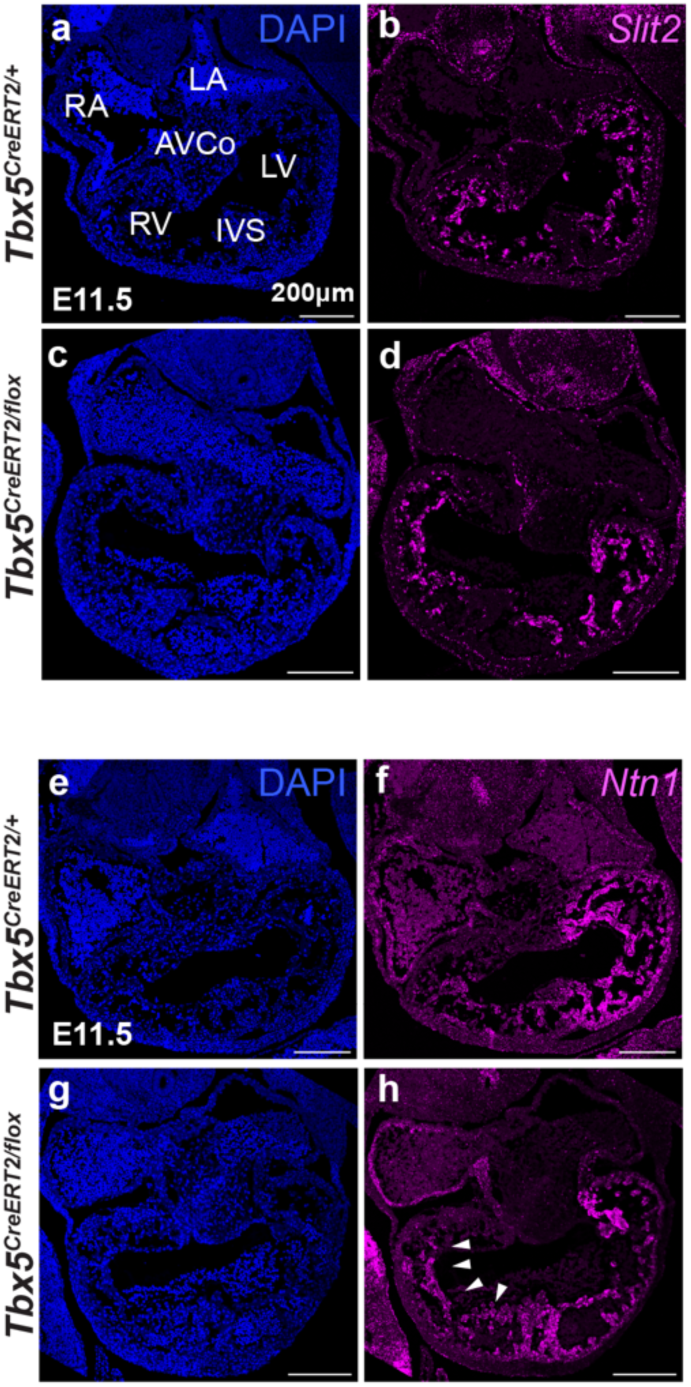
Expression of *Slit2* and *Ntn1* in *Tbx5* mutants at E11.5. Fluorescent *in situ* hybridization of (**a-d**) *Slit2* and (**e-h**) *Ntn1* at E11.5 in *Tbx5^CreERT2/+^* controls (*Tbx5^CreERT2/+^*;*Mef2cAHF-DreERT2*;*ROSA26^Ai6/Ai66^*) and *Tbx5^CreERT2/flox^* mutant (*Tbx5^CreERT2/flox^;Mef2cAHF-DreERT2*;*ROSA26^Ai6/^*^Ai66^) hearts. Arrowheads highlight ectopic expression of *Ntn1* in the RV. All scale bars are 200 microns.

**Extended Data Figure 8.**
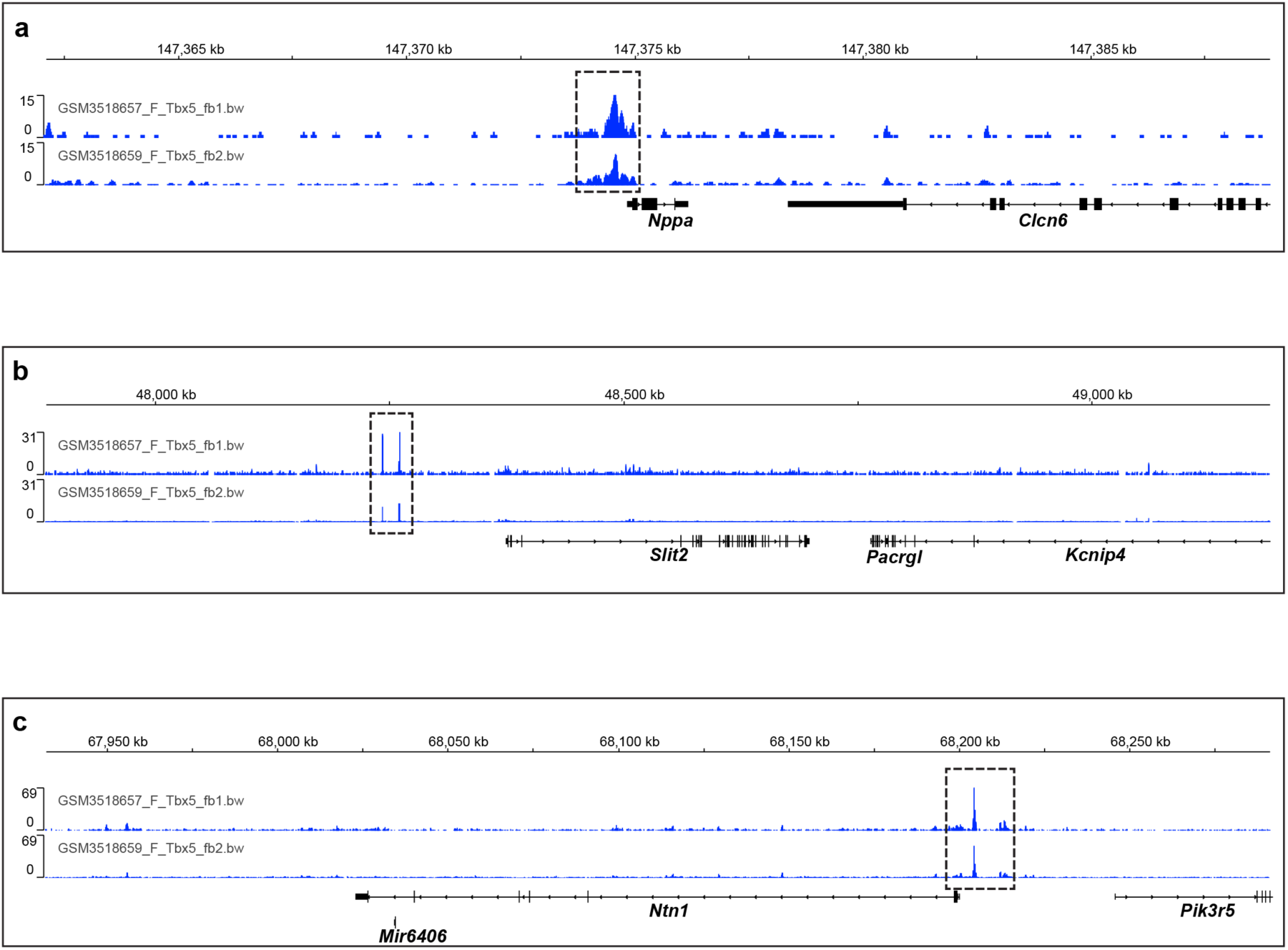
TBX5 chromatin occupancy near *Slit2* and *Ntn1*. **a-c**, Browser tracks of TBX5 ChIP-seq from E12.5 hearts ^55^ show peaks of TBX5 occupancy (dashed black boxes) near promoters of *Nppa*, *Slit2* and *Ntn1*.

**Extended Data Figure 9.**
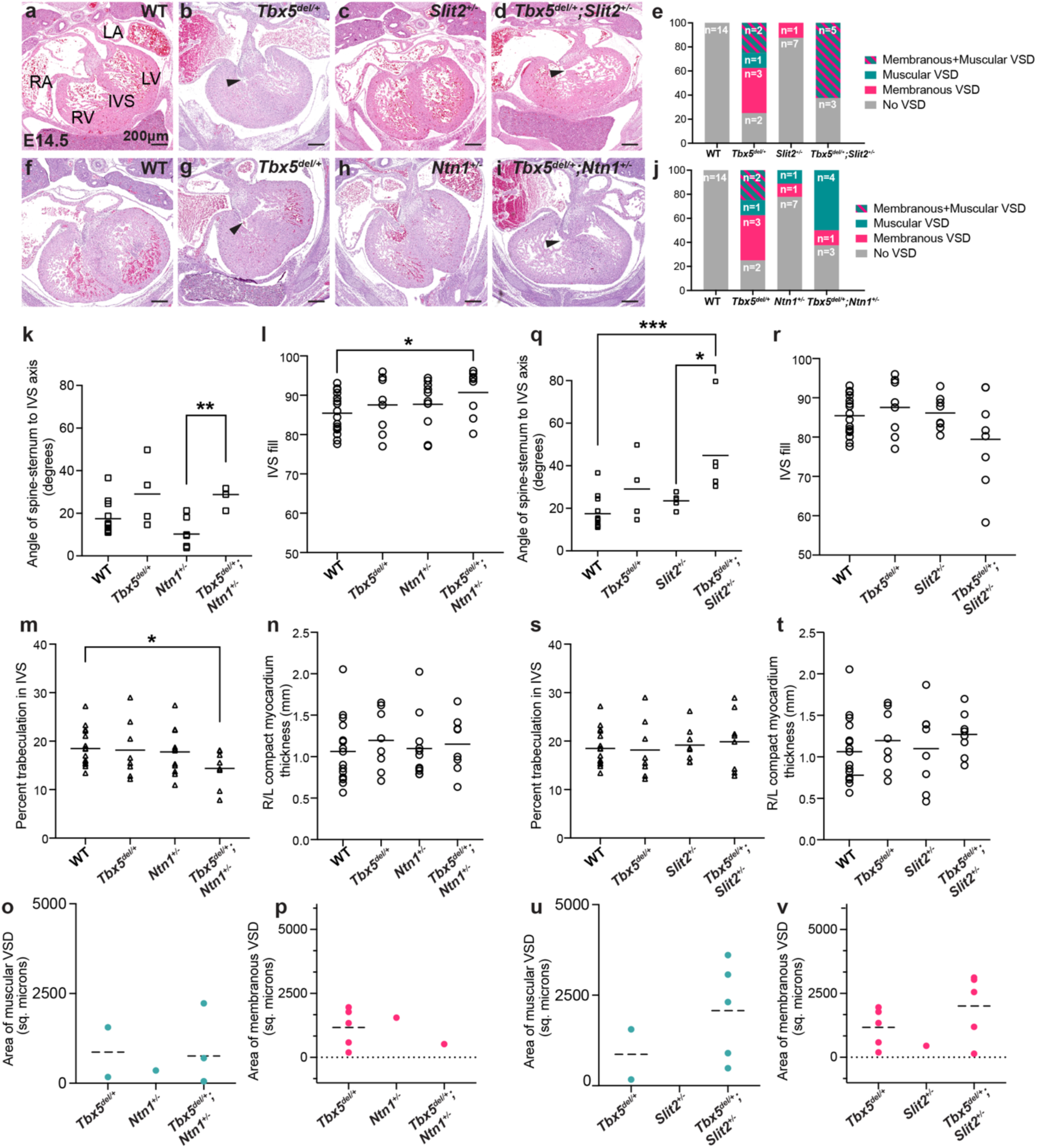
*Tbx5-Ntn1* and *Tbx5-Slit2* genetic interactions. Histology from E14.5 embryos from matings of (**a-d**) *Tbx5^del^*^/+^ and *Slit2*^+/-^ or (**f-i**) *Tbx5^del^*^/+^ and *Ntn1*^+/-^ by genotype. Arrowheads depict muscular ventricular septal defects (VSDs). **e,** Quantification of membranous or muscular VSDs by genotype. *Tbx5^del/+^*;*Slit2^+/-^*compound heterozygous embryos displayed an estimated decrease in the incidence of membranous VSDs, which approached significance (log OR −2.5; p=0.09 in a Generalized Linear Model without adjustments for multiple comparisons). This decrease was relative to the expected incidence of membranous VSDs if there was no genetic interaction between *Tbx5* and *Slit2*. **j**, *Tbx5^del/+^*;*Ntn1^+/-^* compound heterozygous embryos displayed a decrease of membranous VSDs (log OR −5.3; p=1x10^-3^ by a Generalized Linear Model without adjustments for multiple comparisons), relative to the expected incidence of membranous VSDs if there was no genetic interaction between *Tbx5* and *Ntn1,* but not changes to prevalence of muscular VSDs (log OR 0.12; p=0.93). **k-v**, Machine-learning based quantitative morphometry of metrics for (k-p) *Tbx5^del^*^/+^ and *Ntn1*^+/-^ or (q-v) *Tbx5^del^*^/+^ and *Slit2*^+/-^ by genotype. IVS, interventricular septum. Significance determined by two-sided Fisher’s exact test (*p<0.05, **p<0.01, ***p<0.001). (k) **p=0.0059, (l)*p=0.0337, (m) *p=0.0196, (q)***p=0.0008, *p=0.0468. All scale bars are 200 microns.

**Extended Data Figure 10.**
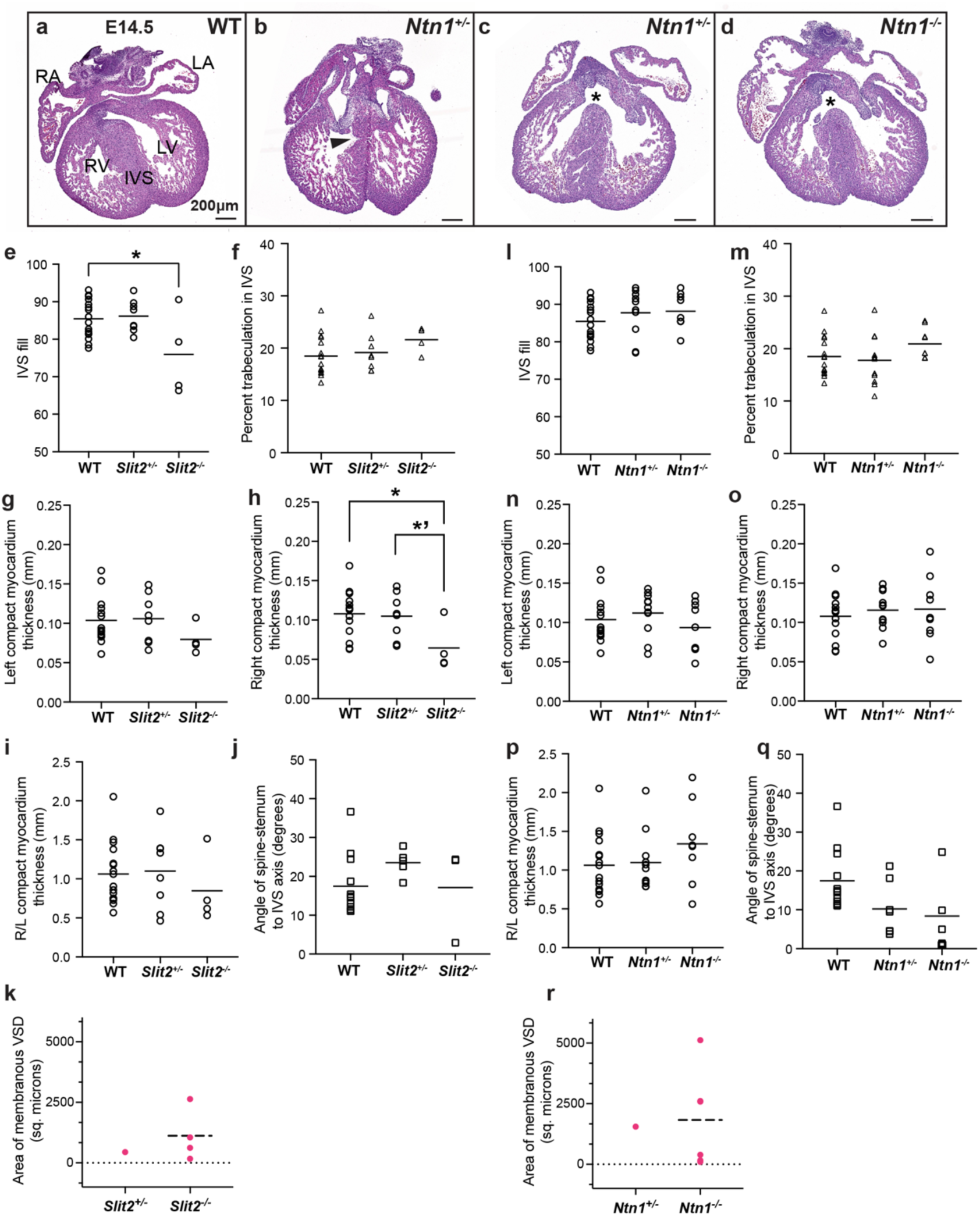
Quantitative morphometry of *Slit2* and *Ntn1* mutant hearts. **a-d**, Additional histology by *Ntn1* genotype at E14.5. Arrowhead, muscular ventricular septal defect (VSD). Asterisk, membranous VSD. Scale bars are 200 microns. **e-p**, Machine learning-based quantitative cardiac morphometrics for (e-k) *Slit2* or (l-r) *Ntn1* mutants. Significance determined by two-sided Fisher’s exact test (*p<0.05). **q, r**, Morphometry of the area of membranous VSDs by *Slit2* or *Ntn1* genotypes. (E) *p=0.0184, (H) *p=0.0216, *’p=0.0492.

